# Molecular architecture of heterochromatin at the nuclear periphery of primary human cells

**DOI:** 10.1101/2025.04.09.647790

**Authors:** Jan Philipp Kreysing, Sergio Cruz-León, Johannes Betz, Carlotta Penzo, Tomáš Majtner, Markus Schreiber, Beata Turoňová, Marina Lusic, Gerhard Hummer, Martin Beck

**Affiliations:** Department of Molecular Sociology, Max Planck Institute of Biophysics, Max-von-Laue-Straße 3, 60438 Frankfurt am Main, Germany; IMPRS on Cellular Biophysics, Max-von-Laue-Straße 3, 60438 Frankfurt am Main, Germany; Department of Theoretical Biophysics, Max Planck Institute of Biophysics, Max-von-Laue-Straße 3, 60438 Frankfurt am Main, Germany; Department of Infectious Diseases, Virology, Heidelberg University, Heidelberg, Germany; Institute of Biophysics, Goethe University Frankfurt, 60438 Frankfurt am Main, Germany; Institute of Biochemistry, Goethe University Frankfurt, 60438 Frankfurt am Main, Germany

## Abstract

In eukaryotes, meters of DNA are packaged into micrometer scale nuclei. Nucleosomes, as the major organizational unit, have been extensively studied in vitro, yet the elaborate 3D structure of chromatin inside cells and its distinct multi-nucleosome arrangements remain poorly resolved. Here, we combine cryo-electron tomography with template matching, subtomogram averaging and molecular simulations to visualize nucleosomes and chromatin structure inside human cells. We confidently assign individual nucleosomes and report their in-situ structure at secondary structure resolution. By tracing linker DNA, we identify multi-nucleosome arrangements and uncover higher-order chromatin structures in situ, including a 37-nm wide, elongated but non-fibrous arrangement. In situ structural biology thus reveals the molecular chromatin organization inside cells and sets the stage for 3D genomics.

## Introduction

The genetic material within each human cell consists of approximately two meters of DNA that needs to be packaged into a nucleus that is six orders of magnitude smaller in diameter. This packaging challenge is compounded by the requirement that the compacted genome must remain accessible for transcription, replication and remodeling in a controlled, efficient, and temporally precise manner. One mechanism by which cells meet this challenge is by wrapping DNA around histone proteins. The resulting protein-DNA complexes form nucleosomes, often depicted as “beads on a string”, which in turn fold into stable, higher-order structures, to finally form the complex three-dimensional arrangement known as chromatin. The hierarchical architecture of chromatin has been studied extensively. Light microscopy imaging and advanced sequencing techniques have revealed chromosome organization, chromatin compartments or topologically associating domains (TADs), while structural biology techniques have been used to analyze the molecular details of how DNA is wrapped around individual reconstituted nucleosomes ^1^. However, how chromatin is organized at the level of multiple nucleosomes inside of cells, in particular how nucleosomes are arranged with respect to each other in native chromatin, remains the subject of active research.

Canonical nucleosomes are formed by a C2-symmetric octamer of two copies of each of the core histones H2A, H2B, H3 and H4. 147 base pairs (bp) of DNA are wrapped in a left-handed superhelical turn around this octamer, and flanked by stiff DNA linkers of variable length that project towards the next nucleosome ^2^. A single H1 linker histone can bind to the DNA adjacent to the core nucleosome superhelix, thereby disrupting the nucleosome’s 2-fold symmetry ^3^. The core nucleosome with H1 and its two DNA linkers, referred to as a chromatosome, is thought to be the building block of higher-order chromatin structures. H1 variants can bind to either of the two DNA linkers, the DNA wound around the histone octamer, or all three DNA structures simultaneously, resulting in differential stabilization and either on-dyad or off-dyad arrangements ^4,5^.

Local chromatin folding is known to be determined by nucleosome organization ^5^. Several models have been proposed for the arrangement of nucleosomes into a 30 nm chromatin fiber, such as e.g. the zig-zag model in which alternating nucleosomes interact, or the solenoid model that implies interactions of neighboring nucleosomes ^5^. Indeed, arrangements of multiple nucleosomes on DNA in vitro are characterized by a zig-zag arrangement, whereby nucleosomes can stack with each other ^6-9^, and the DNA linkers restrain the orientation of consecutive nucleosomes ^10^. In addition, a flat ladder-like structure and twisted arrangements have been described ^5^. Chromatin folding and its compaction behavior is influenced by H1 binding in on-dyad or off-dyad position and on the nucleosome repeat lengths (NRLs) ^11^. Reduced DNA linker lengths and consequently shorter NRLs sterically hinder H1 binding and result in less regular arrays, which in turn are associated with active chromatin ^6,12^. These molecular details have been derived from extensive in vitro structural analyses of reconstituted or purified histones, often using artificial, so-called 601 nucleosome positioning sequence (NPS) repeats ^13^. To which extent these studies reliably recapitulate the architecture of native chromatin in situ, which is not only dependent on NRLs, but also affected by phase separation, ionic strength and various other associated proteins beyond linker H1 ^14,15^, remains largely unknown.

Several recent studies have investigated nucleosome arrangements in different cell types in situ using cryo-electron tomography (cryo-ET) ^16-22^. Although nucleosome stacking was observed ^16,19^, an overall relatively heterogenous chromatin arrangement was reported ^16^. Human cells contained a fibrous chromatin structure lining the nuclear envelope ^18,19^, with exclusion zones beneath nuclear pores ^18^. However, the respective in situ structures of nucleosomes were not resolved to high resolution ^16-19^, and the topology of neighboring nucleosomes was not assessed with respect to the above-discussed structural features that are known to constitute chromatin architecture in vitro. Nevertheless, in vitro and in situ structures combined have pointed to possible higher order structures ^20^.

Here, we determine nucleosome structure and multi-nucleosomes arrangements in situ based on cryo-ET combined with template matching, subtomogram averaging (STA) and molecular simulations. We use resting T cells for our analysis because heterochromatin is prominently enriched in proximity to the nuclear envelope. We resolve the chromatosome to secondary structure in situ; we trace DNA linkers between individual chromatosomes up to kilobase pair (kbp) genomic fragments; we identify key molecular determinants that induce order or disorder in multi-nucleosome arrangements; and we explain how nucleosome stacking as well as orientation and length of the DNA linkers contribute to the ∼37 nm broad, elongated, but non-fibrous arrangement we observe in situ. Our study highlights the potential of in situ structural biology to examine chromatin structure at the multi-nucleosome scale directly within the dense environment of the cell nucleus. As such, it opens the door for 3D genomics and the study of spatial genome regulation within the nuclear space directly inside cells.

## Results

To study multi-nucleosome arrangements in their native state, we vitrified resting T cells from human donors and subjected them to specimen thinning by focused ion beam milling. 14 tilt series were further processed with a projection interpolation algorithm, cryoTIGER ^23^, to improve angular sampling ^23^ reconstructed in novaCTF ^24^ and template matched using GAPSTOP^TM 25,26^ with a nucleosome structure (Fig 1a,b,c) (see Methods and Supplementary Fig 1 for detail). The resulting cross-correlation peaks (Fig 1b) visually coincided with the particles that resembled nucleosomes (Fig 1c). These were considerably more frequent in the nucleoplasm as compared to the cytosol (Supplementary Fig 2), in line with the expected localization of chromatin. We therefore set the threshold for peak extraction at the 99^th^ percentile of cytoplasmic peaks (Fig 1d, see Methods for details). We observed an about 50 nm thick layer densely packed with nucleosomes underneath the inner nuclear membrane (Fig 1a), which is consistent with recent findings on lamin-nucleosome interactions in Murine Embryonic Fibroblasts ^18^.

**Fig 1:**
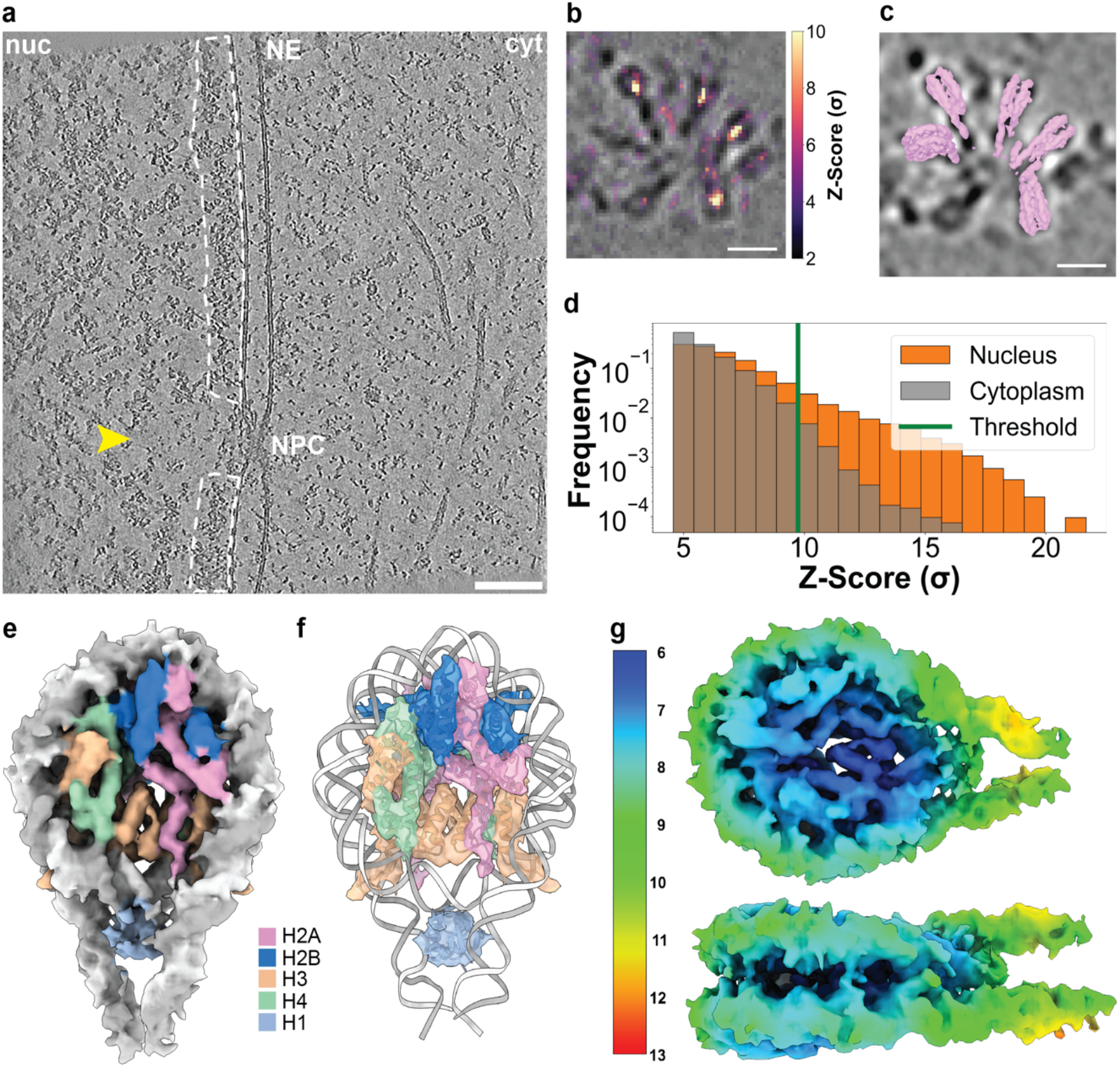
In situ structural analysis identifies nucleosomes inside of cells and resolves their secondary structure. **a** Slice through a cryoCARE-denoised ^28^ tomogram acquired at the nuclear periphery of resting T cells. Chromatin-free space is indicated with a yellow arrow, a dense layer of chromatin underlying the nuclear envelope is framed white. **b** A local feature of multiple nucleosomes superimposed with the respective cross-correlation function obtained by template matching. **c** The same local feature but with the identified nucleosomes shown as isosurfaces. **d** Histogram of Z-scores for the cytosolic and nucleoplasmic region for exemplary tomogram. **e**,**f** STA map of the human chromatosome, color-coded by DNA strands (white and gray) and histone proteins H2A (pink), H2B (blue), H3 (amber), H4 (green) and H1 (light blue). The atomic model of the canonical chromatosome (PDB: 7DBP ^27^) was fitted into our map. **g** Color-coded local resolution STA map of the human chromatosome (from high resolution in blue to low resolution in red, in Å). Scale bars 100 nm in **a** and 10 nm in **b**,**c**.

STA (performed with non-interpolated data) confirmed that the extracted peaks primarily account for nucleosomes. We obtained an in situ structure of the chromatosome from resting T cells locally resolved to 6.4 Å (7.3 Å overall) with imposed C2 symmetry (Fig 1e-g). It closely resembles previous in vitro structures ^27^ of canonical chromatosomes with H1 bound and two DNA linker arms. The STA map clearly showed the secondary structure features of all core histones, the major and minor groove of the wrapped DNA emerging into the DNA linkers, and the histone tails of histone H3 and H4 projecting outwards through the DNA. Without imposed 2-fold symmetry the resolution was slightly reduced (Supplementary Fig 1, 3, Supplementary Table 1), but the asymmetrically associated histone H1 more clearly resolved. Extensive classification attempts also identified nucleosomes in which H1 was shifted to either DNA linker, characteristic for off-dyad binding (Supplementary Fig 4). These data suggest that an H1 histone variant, which interacts with both the linker DNA arms and the DNA superhelix at the dyad, plays a role in restraining the positioning of linker DNA. This regulation influences the arrangement of nucleosomes relative to one another and is particularly abundant at the nuclear periphery in resting T cells. In addition, to the best of our knowledge, our nucleosome structure from resting T cells represents the smallest non-oligomeric structure resolved to secondary structure resolution by STA in situ to date, with a molecular weight of about 250 kDa.

We next used the set of detected nucleosomes to determine chromatin density (Fig 2). Clustering analysis revealed an overall densely packed nucleoplasm with some occasional chromatin-free patches, e.g., underneath nuclear pores (Fig. 1a, 2a,b). Based on the identified nucleosomes, we determined the local density distribution by splitting the nuclear volume into non-overlapping 50-nm cubes and counting the number of nucleosomes within. The analysis revealed that the local nucleosome concentration can be very high, exceeding 200 mg/mL, a value previously reported in the literature ^29^ (Fig 2a,e). We also quantified how much nuclear volume was occupied by the nucleosomes as compared to chromatin free space (Fig 2b-d), and found that less than half of the volume at the nuclear periphery is chromatin free. Finally, we quantified native chromatin in terms of the nucleosome pair distribution function (Fig 2f). We find chromatin to be intermediate between a liquid and gas with two short-range peaks at 7 and 11 nm center-to-center distance likely representing stacked (Supplementary Fig 8) and edge-contacting nucleosomes, and a gradual decay beyond 20 nm indicating a weak tendency to condense (Fig 2f). The relatively featureless curve provides no evidence for long-range order in chromatin, arguing against fibrous chromatin structures.

**Fig 2:**
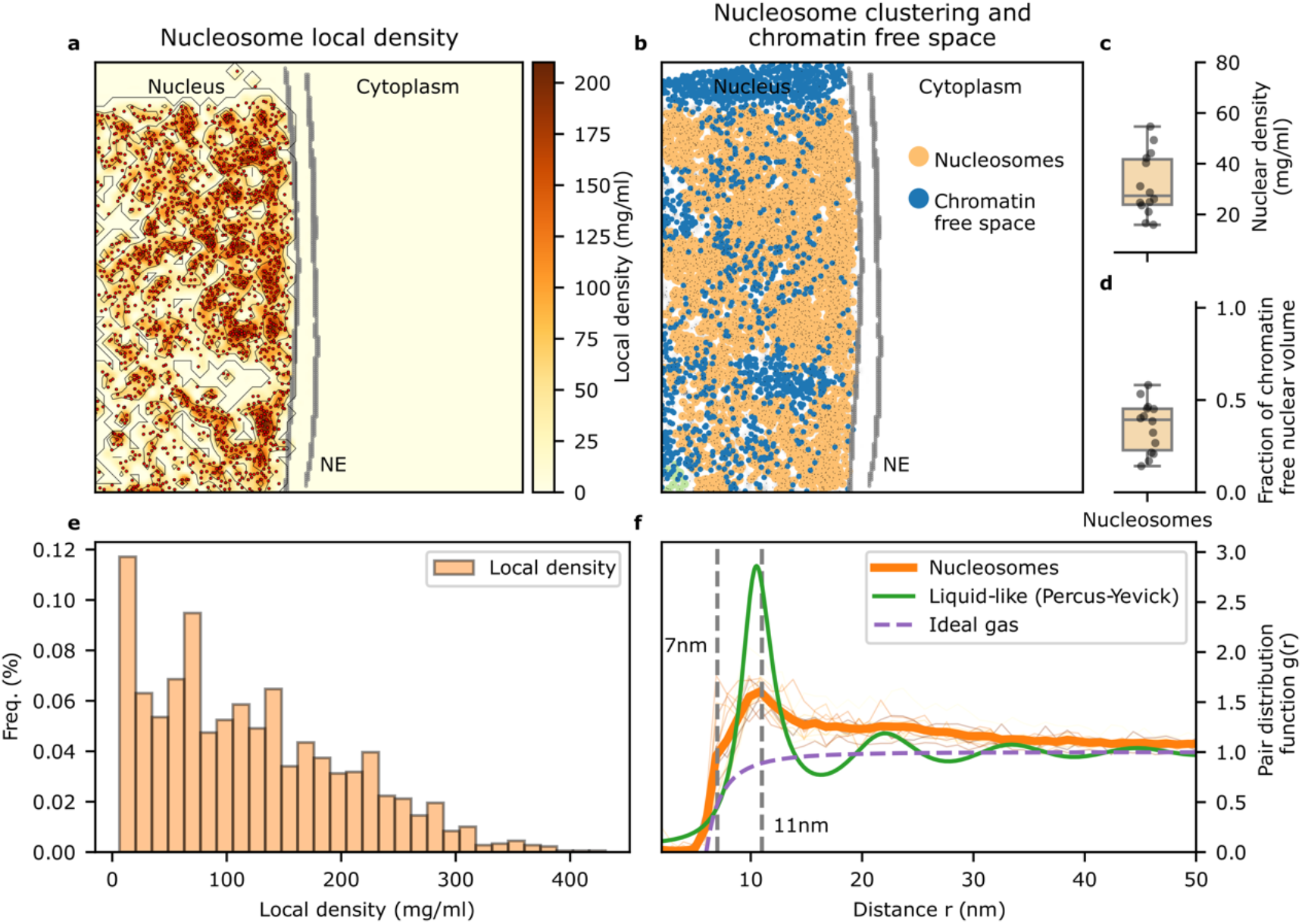
Spatial properties of heterochromatin at nuclear periphery of T cells. **a** Nucleosomes local density, **b** hierarchical density-based clustering (orange and green clusters) and chromatin free space visualized from the same exemplifying tomogram as in Fig 1a. **c** Overall nucleosome density, **d** fraction of chromatin free nuclear volume, **e** local density distribution over all tomograms and **f** pair distribution function of the nucleosomes. Reference curves for the ideal gas and liquid-like distributions ^30^ are included for comparison.

To analyze multi-nucleosome arrangements in higher detail, we connected consecutive nucleosomes by tracing the DNA linkers. We first used a distance-based clustering as implemented in cryoCAT ^31^ to identify the candidate nucleosomes to which they could be connected by a DNA linker. For this, we select neighbors that have at least one of their linker arms (linker 1 and linker 2 in Supplementary Fig 5b) within 20 nm of at least one of the two linker arms of the other nucleosome. Within a given local cluster defined by this distance criterion, the positions and orientations allowed us to connect candidate nucleosome pairs by using a physics-based DNA model of the bending energy *U*_bend_ of the DNA linker (Fig 3a and Supplementary Fig 5a). This bending energy *U*_bend_ is calculated based on the linker length (L), the bending angle (θ), and the configuration of the DNA linker between a pair of nucleosomes that is connected (where Γ denotes the specific linker connection in Supplementary Fig 5b) (see Methods for details) as follows:

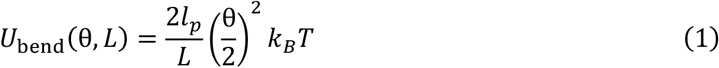

**Fig 3:**
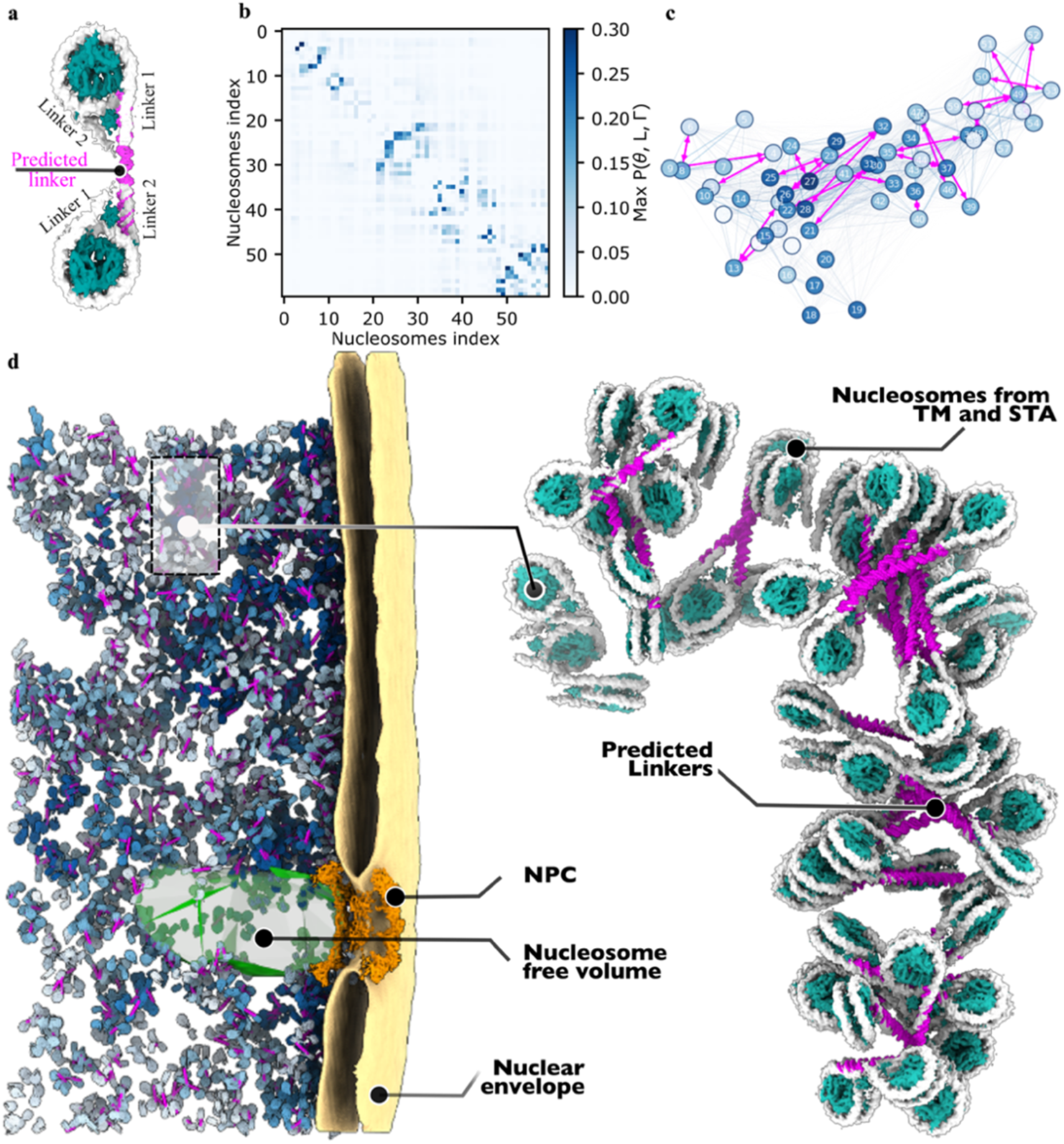
Cryo-ET and physics-based model allow kbp-long chromatin to be visualized. **a-c** A physics-based model of nucleosome pairs is used to add DNA linkers (magenta) to in situ nucleosome DNA (white) (a). All four distinct possibilities to connect two nucleosomes with two linkers each are scored, resulting in connectivity matrix for all nucleosomes within a given cluster (see Methods and Supplementary Fig 5 for an example) (b). The connectivity matrix from **b** is represented as graph, with nucleosomes as nodes connections as edges (c). Nodes are projected on the x-y plane and colored according to the z-coordinate. Edge thickness and color are proportional to the connection probability. Assigned linkers are shown as solid arrows between nodes. **d** Visualization of an exemplifying tomogram (corresponding to Fig 1a) showing the densely interconnected chromatin structure within the nucleus. Nucleosomes are colored according to the length of the traced nucleosome chain, from short (white-light blue) to long (dark blue); predicted linkers are shown in magenta. The nuclear envelope membrane was segmented with MemBrain-seg ^32^.The zoom-in is the 3D representation of 60 traced nucleosomes in b-c.

Here *l*_*p*_ ≈ 50 nm is the B-DNA persistence length at physiological conditions, *k*_*B*_*T* is the thermal energy and *L* = *θr* is the length of the arc of a circle sector of radius *r* and central angle *θ* (in radians). The radius *r* is related to the Euclidian distance *D* between the ends of the linker arms on the two nucleosomes (Supplementary Fig 5a) through

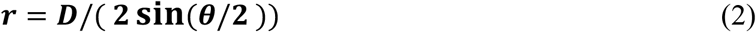

With the energy function in equation 1, we estimated the probability that the DNA linker arms of nearby nucleosomes are connected by a DNA linker as

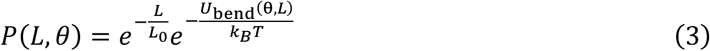

where *L*_0_ is a reference linker length, which we set to *L*_0_ = 15 nm as an expected typical linker length. The resulting probability function thus penalizes very long but straight as well as very short but strongly bent linkers. Both *L*_0_ and *l*_*p*_ are parameters of the algorithm that can be adjusted according to species, tissue, cell type, cell state, and genome location, if needed.

In a spatial cluster of *N* nucleosomes, we estimated in this way the relative probabilities for each of the N × *N* × 4 possible connections to be formed, where the factor of 4 accounts for the four possible connections of the two arms on each nucleosome. The resulting connectivity “matrix” *P*_*ijk*_ = *P*(*L*_*ijk*_, *θ*_*ijk*_) of size N × *N* × 4 accounts for all possible connections in a probabilistic manner, where *i* and *j* index the *N* nucleosomes of the cluster and *k* indexes the four possible linker-arm connections between two nucleosomes. Connections are formed in a step-wise manner whereby the most likely connections are assigned in a “greedy” algorithm. This algorithm has three steps that are applied iteratively:

1. Identify the highest value in the current *P*_*ijk*_ matrix, which gives a connection (*n, m, l*) between nucleosomes *n* and *m* for linker connection *l*
2. If the probability exceeds a predefined threshold, *P*_*nml*_ > *P*_*min*_, add (*n, m, l*) to the list of connections; otherwise stop the search
3. If (*n, m, l*) has been added as new connection, update the connectivity matrix by setting *P*_*nml*_ to zero; in addition, we set to zero all entries in the *P*_*ijk*_ matrix that lead to an unfeasible solution with a closed form of the DNA (i.e., with the DNA emanating from one nucleosome arm and eventually returning to the other arm of the same nucleosome if traced along the list of connections) or to multiple connections to the same linker arm
4. Return to step 1

Results for the matrix *P*_*ijk*_ = *P*(*L*_*ijk*_, *θ*_*ijk*_) of relative connection probabilities between nucleosome arms are visually illustrated in Fig 3b and Supplementary Fig 5c. Shown is the 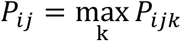 . We find that multiple nucleosomes can be confidently linked by choosing pairs connected with high probability. Similar probabilities for multiple possible pathways rarely arise, leaving the linker solution clear in the vast majority of assigned nucleosome pairs (Supplementary Fig 6). Therefore, a simple “greedy” algorithm sufficed here to connect nucleosomes, starting with the pair of nucleosome arms connected with the highest probability to the not-yet-connected pair with the next-highest probability and so on, always ensuring that no closed DNA loops are formed. The algorithm stops if the next connection falls below a preset threshold, here set to *P*_*min*_ = 0.1. Although most nucleosome connections are pairs (Fig. Supplementary Fig 7), in some cases, up to kbp-long chromatin fragments become interconnected (Figs. 3c,d, and Supplementary Fig 7), and ultimately break as the chain of nucleosomes reaches the edge of the FIB lamella.

To validate the outcome of the algorithm, i.e. the assigned nucleosome pairs, we subjected the predicted DNA linkers to STA. The averages clearly showed elongated density with the expected width underscoring that we had identified regions that contained DNA (Supplementary Fig 6). We next grouped the subtomograms according to the predicted linker length, and subjected the individual classes to STA separately. We found that the length observed in the averages corresponds to the linker length predicted by our algorithm (Supplementary Fig 6).

The connected chromatin fragments provide a close-up view of 3D heterochromatin organization. In the following, we will refer to consecutive nucleosomes as i, i+1, … , i+n species (Fig 4a), as seen in an exemplifying tetra-nucleosome arrangement from our tomograms (Fig 4b). This data allows us to systematically determine center to center distance of consecutive nucleosomes (Fig 4a) and the DNA linker length (Fig 4b,c). The average center-to-center distance between consecutive nucleosomes, i and i+1, was 26.2 nm, corresponding to a ∼37 nm edge to edge boundary. This corresponds to nucleosome repeat lengths (NRLs) of ∼200 bp (Fig 4c). Further, we systematically quantified several parameters, including the i, i+1 distance r_i,i+1_ (Fig 4a,d), i, i+2 distance r_i,i+2_ (Fig 4a,e) and the i,i+1,i+2 angle O_i_ (Fig 4a,f). A prominent organizational feature is the stacking of the i and i+2 nucleosome at an average center-to-center distance of 11 nm (Fig 4e), whereby the connecting i+1 nucleosome is in an opposing position. This is exemplified by nucleosomes 1-2-3 in Fig 4b and g. STA confirmed two different types of stackings, with i and i+2 nucleosomes either directly on top of each other or in a laterally shifted position (Supplementary Fig 8). The center-to-center distance distribution of the i, i+2 nucleosomes shows a second, even more prominent peak at an average distance of 28 nm, that is representative for non-stacking arrangements. This is exemplified by nucleosomes 2-3-4 in Fig 4b and g.

**Fig 4:**
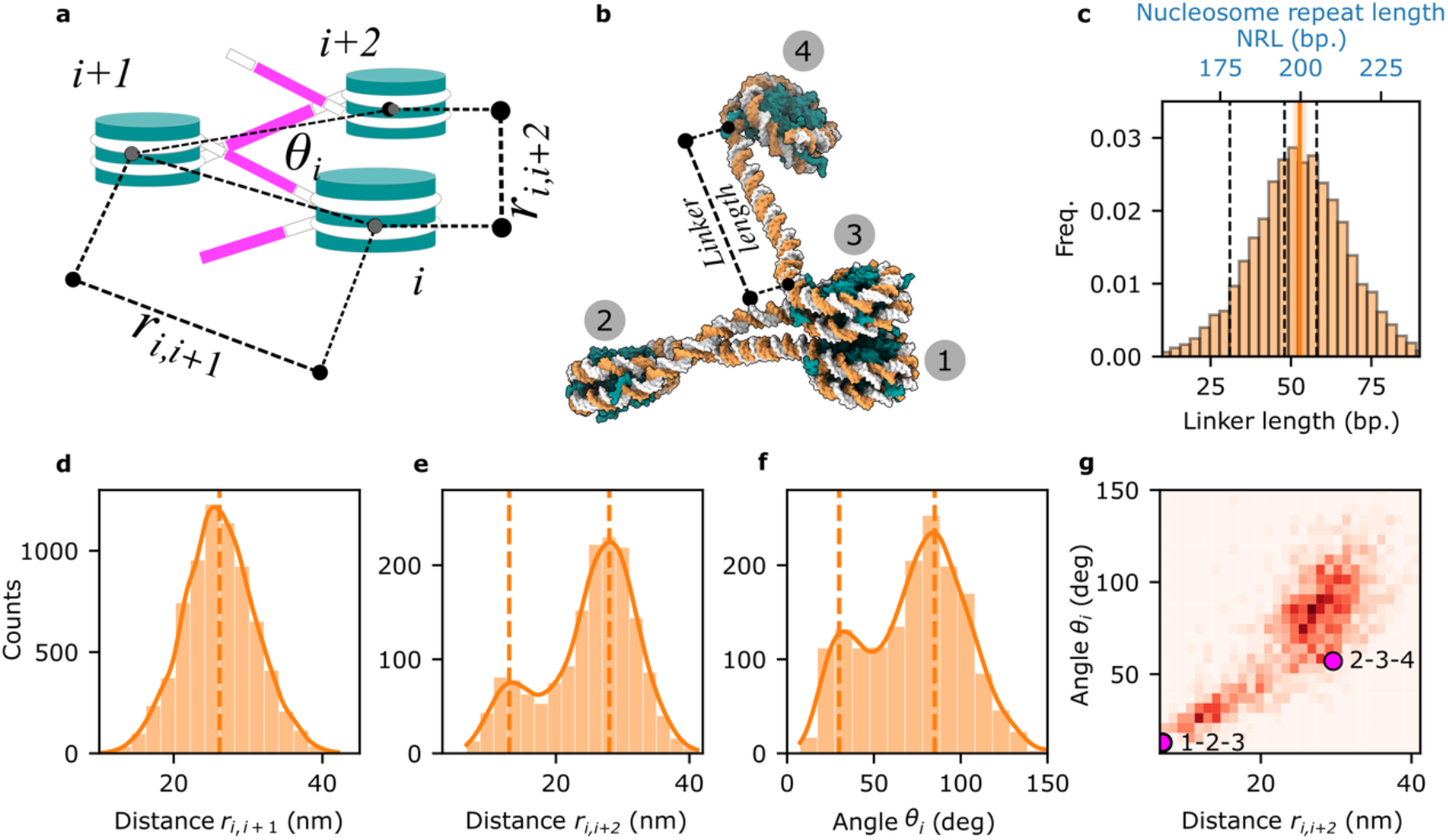
Molecular determinants of multi-nucleosomes that define heterochromatin architecture in situ. Panels **a-g** depict the multi-nucleosome arrangement analysis described by the center-to-center distance; **c** DNA linker length distributions, with dashed vertical lines for chromatin marks taken from ^12^: heterochromatin (205 bp), euchromatic gene bodies (195 bp), active promoters and enhancer (∼178 bp); **d** the distance between connected nucleosomes r_i,i+1_; **e** the second-neighbor distance r_i,i+2_; **f** the angle θ_*i*_ formed by the connected nucleosomes i, i+1 and i+2 (refer to the cartoon in the upper left); **g** the correlation between r_i,i+2_ and θ_*i*_.

The distribution of angles formed by three consecutive nucleosomes is bimodal with peaks at about 30 and 85 degrees (Fig 4f,g). An angle of about 30 degrees is observed when the i and i+2 nucleosome stack with each other, turning to 85 degrees without stacking. These two alternative tri-nucleosome arrangements are frequently re-occurring motifs, and often observed consecutively, as exemplified in (Fig 4b). In a plot of the i, i+2 nucleosome distance against the respective angle, which has been previously used to visualize the molecular determinants of chromatin folding in vitro ^6^, two clusters are apparent (Fig 4g). The two clusters are associated with preferred angles, given linker lengths, and stacking (Fig 4b). Strikingly, these two clusters have been previously observed for tetra-nucleosomes in vitro ^6^, showing that those constituting features of chromatin architecture can be reconstituted in simplified systems. This analysis suggests that the above-described parameters are the key molecular determinants of chromatin folding in vivo.

To explore the dynamics of those key molecular determinants, we selected a representative patch, modeled it in atomic detail and subjected it to all-atom molecular dynamics simulations (Fig 5a). The overall molecular arrangement was stable throughout the simulated 200 ns, indicating a reasonable structural representation of the highly charged DNA and histones. The largest motion occurred at a terminal, singly connected nucleosome, which tightened its stacking interaction. The simulation illustrates the overall dynamic, but nonetheless well-defined ∼37 nm wide arrangement. We observe changes in the angle of a tri-nucleosome of about 10° across the trajectory (Figs. 4f, 5b). The histone tails were initially modeled in random conformations (see Methods for details). As expected from their sequence rich in basic residues, they engage in multiple contacts with neighboring nucleosomes and linker DNA during the simulation (Supplementary Movie 3). Interestingly, histone tails also make contact with nucleosomes that are not directly proximate in the linear DNA chain, illustrating the multivalency of the system. In particular, we observe the H1 tails binding to distant nucleosome cores and linker DNAs, promoting chromatin compaction. Furthermore, the formation of nucleosome stacking interactions is observed (Fig 5c), partially mediated by H4-tail interaction with the histones of the neighboring nucleosome (Fig 5c) as has been previously reported ^33^. Under physiological conditions, post-translational modifications on histone tails may reduce the charge and thus weaken those interactions.

**Fig 5.**
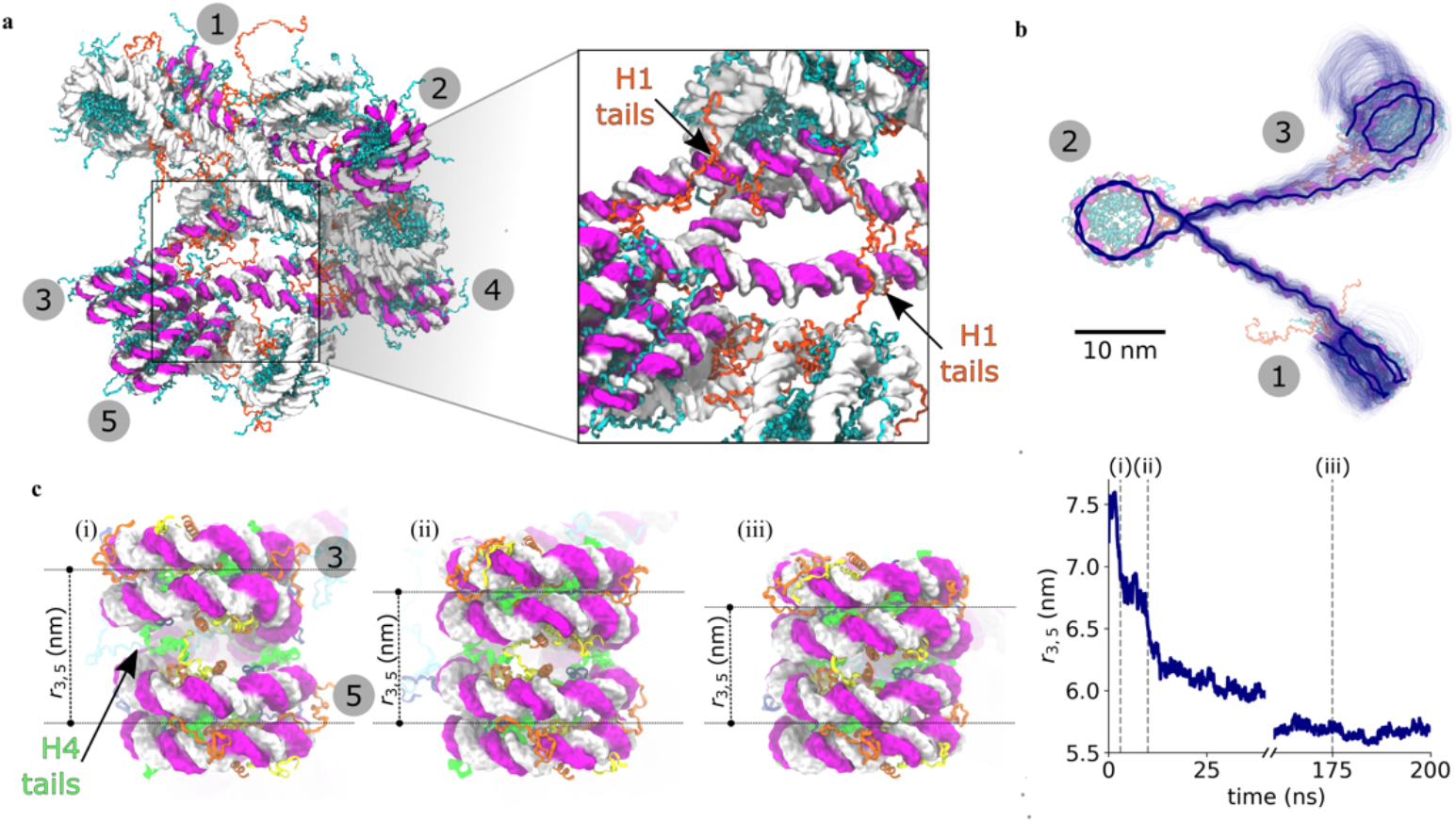
Molecular dynamics (MD) simulation of 13 chromatosomes, 5 of which are connected. **a** exemplifying penta-nucleosome arrangement obtained from a tomogram and modeled in atomic detail, inset highlighting histone H1 tails. **b** visualization of the dynamics of angle formed by a tri-nucleosome arrangement. **c** formation of a nucleosome stacking interaction initially mediated by the H4 tail in (i), progression from (i) to (iii).

## Discussion

We have demonstrated that nucleosomes can be confidently identified in cryo electron tomograms and that the trajectory of their DNA linkers can be estimated. Thus, our study demonstrates that the analysis of multi-nucleosome arrangements is feasible in situ, and allows us to extract the key molecular determinants of higher-order heterochromatin structure. Overall, our analysis points to the following molecular traits as determinants: DNA linker lengths or NRLs, frequent nucleosome stacking, angular restraining by relatively inflexible DNA linkers and H1 binding, multi-valent contacts of histone tails to neighboring histones that must not necessarily be close in sequence. Thereby some features, such as smaller NRL variability, smaller tri-nucleosome angles and angular restraining by H1 binding, promote nucleosome stacking and thus induce order. Conversely, larger NRL variability and larger tri-nucleosome angles will counteract nucleosome stacking, and thus induce disorder.

A regular 30 nm fiber has been suggested to be a major feature of heterochromatin in situ, but remains debated ^3,7,8,20,34,35^. Whereas an organized 30-nm fiber is absent, we find a more heterogeneous, but still elongated structure with a characteristic width of ∼37 nm. Differences to the aforementioned 30 nm fiber are that the DNA linker length observed in this study was on average ∼17.5 nm, while a 30 nm fiber would require considerably shorter linker DNA. Furthermore, deviations in linker length, lateral displacement of stacked nucleosomes and occasionally occurring unusual angles break with the regularity of chromatin folding in situ. This results in a more heterogenous arrangement that is about 37 nm broad and elongated, thus appearing somewhat fibrous without showing the hallmark features of a highly ordered material (Fig 2f).

We focused our analysis on the nuclear periphery, where heterochromatin ^36^ is known to attach to the inner nuclear membrane ^37,38^. Therefore, it is likely that transcriptionally inactive chromatin is prominently contained in our data. Traditional imaging techniques such as light microscopy or plastic embedding electron microscopy suggest that the heterochromatin layer underneath the nuclear envelope is variable in thickness, but may considerably extend into the nucleus, such that active chromatin may be outside our present field of view. This would explain why our extensive efforts to template match polymerases and chromatin remodelers, did not yield any statistically compelling results. Interestingly, we did observe a layer of chromatin that was firmly attached to the nuclear envelope (Fig 1a,b). This morphologically distinct layer was only about 50 nm thick, and is thus beyond the diffraction limit for light microscopy.

Our in situ structural analysis can be compared directly to genomics data. We quantified DNA linker length in our tomograms, finding on average ∼200 bp separating the 147 bp wrapped around the nucleosome octamers in resting T cells (Fig 4c). While our data only capture the nuclear periphery, genomics methods were previously used to estimate nucleosome spacing in T cells in a genome-wide and histone-mark specific manner ^12^. More specifically, spacing for chromatin marks H3K27me3, H3K9me3 attributed to heterochromatin (205 bp) and H3K27me1 ^39,40^ attributed to euchromatic gene bodies (195 bp) was higher in comparison to H3K4me1 and H3K27ac (∼178 bp) that were attributed to enhancers and super enhancers ^41^. The distribution in our data considerably overlaps with the values expected for heterochromatin (Fig 4c).

In addition, we hereby demonstrate the power of cryo-ET to analyze smaller complexes in the range of only about 250 kDa at subnanometer resolution. This may have broader applications and enable the in situ structural analysis of a range of biological processes. Regarding chromatin, further technical improvements in tomographic data acquisition and processing schemes should yield cryo electron tomograms of active chromatin in the near future, thus opening up the exciting possibility of observing gene expression at work. Such data may also allow pinpointing histone variants by extensive image classification combined with molecular simulations, and to spatially associate them with proximate regulatory complexes. Thus, genomics and in situ structural biology may go hand in hand in the future towards improving our understanding of chromatin structure and function relationships. Taken together, our study bridges our understanding of chromatin from molecular to multi-nucleosome arrangements and illustrates the potential of in situ structural biology in contributing to future mechanistic analyses of chromatin compaction and remodeling.

## Methods

### Preparation of primary CD4^+^ T cells

Detailed CD4+ T cells purification method used is described in Lucic et al., 2019. Briefly, cells were obtained by using RosetteSep Human CD4^+^T cells enrichment cocktail (STEMCELL technologies) and isolated by Ficoll gradient centrifugation. Following the lysis of red blood cells, purified T cells were cultured in RPMI-1640 10% FBS and penicillin/streptomycin, in the presence of IL2 10 ng/uL and incubated at 37°C in a 5% CO_2_ atmosphere. Cells were then transferred to 20mM Hepes pH 7.9 RPMI-1640 at a concentration of 4 × 10^6^ cells for further use.

### Cell vitrification and cryo-FIB milling

3.5 µL of cell suspension of resting primary human CD4^+^ T cells was applied on glow discharged 200-mM mesh EM gold grids (Quantifoil Micro Tool GmbH) coated with R 2/2 holey SiO2 films and plunge frozen in liquid ethane –183°C using a Leica EM GP2 grid plunger (Leica Microsystems) with the blotting chamber at 37°C. The grids were blotted from the back (cell-free) side for 4-5 seconds with a Whatman filter paper, Grade 1 before plunge freezing.

The cells were then cryo-FIB milled using an Aquilos 2 microscope (Thermo Fisher Scientific) in a similar fashion to a previously described workflow ^42^. In brief, samples were coated with an organometallic platinum layer for 10 sec and additionally sputter coated with platinum at 1 kV and 10 mA current for 10 sec. The lamella milling was performed with AutoTEM (version 2.4.2) (Thermo Fisher Scientific) in a stepwise manner with a 30kV ion beam while reducing the current from 1000 pA to 50 pA. Final polishing was performed with 30 pA current with a lamellae target thickness of 200 nm.

### Cryo-ET data acquisition

Cryo-ET data for resting T-cells were collected at 300 kV on a Titan Krios G4 microscope (Thermo Fisher Scientific) equipped with a E-CFEG, Falcon 4 direct electron detector (Thermo Fisher Scientific) operated in counting mode and Selectris X imaging filter (Thermo Fisher Scientific). Montaged grid overviews were acquired for each grid with a 205 nm pixel size. Individual lamellae montages were taken with 3 nm pixel size. Tilt series were acquired using SerialEM (version 4.0.20) ^43^ in low dose mode as 4K x 4K movies of 10 frames each and on-the-fly motion-corrected in SerialEM. The magnification for projection images of 64000x corresponded to a nominal pixel size of 1.971 Å. Tilt series acquisition was started from the lamella pretilt of ± 8° and a dose-symmetric acquisition scheme ^44^ with 2° increments grouped by 2 was applied, resulting in 61 projections per tilt series with a constant exposure time and targeted total dose of ∼135 e^-^ per Å^2^. The energy slit width was set to 10 eV and the nominal defocus was varied between -1.75 to -4.25 µm. Dose rate on the detector was targeted to be ∼5 e^-^ /px/sec.

### Tomogram reconstruction

Dose exposure correction for the motion corrected tilt series was performed as previously described ^45^ using a Matlab implementation that was adapted for tomographic tilt series ^46^. Poor quality projection images were removed after visual inspection. The dose-filtered tilt series were then aligned with patch-tracking in AreTomo (v2.0) ^47^ and reconstructed as back-projected tomograms with SIRT-like filtering of 15 iterations at a binned pixel size of 7.884 Å (bin4) in IMOD (version 4.11.5) ^48^. The 14 tomograms containing a nuclear envelope were selected for further processing. These tomograms were then denoised with the cryoCARE software package ^28^ for improved visualization.

For the same tomograms, reconstruction with 3D-CTF correction was also performed using novaCTF ^24^ with phase-flip correction, astigmatism correction and 15 nm slab. Tomograms were binned 2x and 4x using Fourier3D ^49^. For compatibility with Relion 3.1 ^50^ and M ^51^ these 14 tilt series were reprocessed in Warp ^52^ with the alignment obtained from AreTomo ^47^.

### Nucleosome template matching, including initial template generation

To obtain initial particle coordinates and orientations, the bin2 (3.942 Å/px) 3D-CTF corrected tomograms were template matched with an existing nucleosome structure (EMD-3947) (lowpass filtered to 13 Å) and 5 degree global angular sampling in GAPSTOP^TM 25,26^. The resulting constrained cross correlation (CCC) volumes were used to extract positions and coordinates of peaks with a CCC score 4.5 sigmas above the mean CCC score value. The candidate particles determined through template matching were extracted as subtomograms in Warp (v1.09) ^52^ at bin2. Subsequent junk classification and refinement in Relion (v3.1) ^50^ lead to an initial STA nucleosome structure that was then used as an improved search template for further template matching.

To improve template matching performance, we used the cryoTIGER computational workflow ^23^ to interpolate between tilt stacks using the FILM algorithm ^53^, as detailed in ^23^. The aligned bin2 tilt stacks were used as inputs for cryoTIGER where one tilt image was interpolated in between two real tilt images for each stack. The combined stacks containing real and interpolated tilt images were used for novaCTF ^24^ reconstruction using the same parameters as above. The resulting tomograms were template matched with the generated initial STA nucleosome structure without DNA linker arms (lowpass filtered to 13 Å) and 5 degree global angular sampling in GAPSTOP^TM 25,26^. Since all tomograms contained both cytoplasm and nucleoplasm the two regions were manually segmented in IMOD ^48^ for each tomogram. The constrained cross-correlation (CCC) peaks in both regions of the tomogram could thereby be compared and a peak extraction threshold for nucleoplasmic peaks could be set per tomogram. This threshold corresponded to the 99^th^ percentile of CCC values for cytoplasmic peaks in each tomogram (Fig 1d), allowing for extraction of a high-confidence nucleosome particle list. This particle list was further cleaned by excluding clashing particles with a nucleosome shape mask around each peak in cryoCAT ^31^

### Nucleosome subtomogram averaging

The 33,560 particles determined through template matching were extracted as bin1 (1.971 Å/px) subtomograms in Warp ^52^ (no interpolation between tilt stacks) and subjected to subtomogram averaging and alignment in Relion 3.1 ^50^. Multiple attempts at classifying out junk particles were made, all resulting in no significant percentage of particles being classified as junk. The particle set was then imported into M ^51^ to perform multi-particle refinement of the tilt-series and the nucleosome. Geometric and CTF parameters were refined in a sequential manner. Afterwards, new M-corrected bin1 (1.971 Å/px) subtomograms were again refined in Relion 3.1. This resulted in a chromatosome structure resolved to 7.7 Å with C1 symmetry and a 7.3 Å chromatosome structure with C2 symmetry imposed (at FSC 0.143 cutoff). Locally the chromatosome with C2 symmetry imposed was resolved up to 6.4 Å. The particles refined with C1 symmetry were furthermore classified based on potential stacking neighbors and around the H1 linker histone binding site.

### Nucleosome Analysis

To determine the organization, spatial properties and connectivity of heterochromatin at the nuclear periphery inside T-cells, we used the positions obtained from subtomogram averaging and M refinement for further spatial analysis.

### Clustering Analysis

To determine how nucleosomes group together within heterochromatin, we performed cluster analysis of the positions of the nucleosomes using the hierarchical density-based spatial clustering of applications with noise (HDBSCAN) ^54^ clustering algorithm, with a minimum cluster size of 7. HDBSCAN is a density-based clustering algorithm suitable for data with variable densities. For the visualization, each identified cluster was plotted in a unique color to distinguish it from others (Fig 2a).

### Nucleosome density

Nucleosome mass density was estimated for each tomogram using two different methods. First, we determined the average nucleosome density in the nucleus by calculating the ratio of the chromatosome mass ^27^ (242 kDa = ^-^ 4.018507×10^-19^ grams each) to the volume of the convex hull surrounding the particles. The convex hull was estimated using *scipy*.*spatial*.*ConvexHull* ^55^. We also estimated the local mass density of nucleosomes by counting nucleosomes in a cubic grid of 50 nm edge length.

### Fraction of the chromatin-free nuclear volume

To further quantify heterochromatin compaction, we used Monte Carlo integration to estimate the fraction of the nuclear volume that is chromatin-free. First, we defined the volume of the nucleoplasm by manually segmenting the nucleus in each tomogram using IMOD ^48^. We estimated the nuclear volume by calculating the volume of the convex hull that surrounding the segmented mask.

We then attempted to draw 10,000 random particles within the convex hull of the nucleus, with a radius of 5 nm. To determine the relative volume occupied by nucleosomes, we repeated this process taking into account the positions of nucleosomes, which were modeled as spheres of 5 nm radius. A particle was accepted only if it did not collide with more than two nucleosomes. The latter was used to account for isolated nucleosomes. Finally, we calculated the ratio of accepted particles from the nucleosome-inclusive generation to the total number of random particles generated within the nucleoplasm.

### Pair distribution function

To quantify the spatial organization and analyze the local structure of nucleosomes, we calculated the normalized pair distribution function, *g*(*r*) per tomogram. This function quantifies the probability of finding a pair of nucleosomes separated by a distance *r* and allows us to determine the degree of spatial ordering or clustering of nucleosomes.

The normalized pair distribution function, *g*(*r*), was calculated as the ratio:

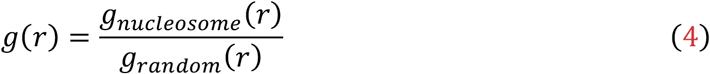

where *g*_*nucleosome*_ (*r*) is the pair distribution function obtained from a histogram of nucleosome positions, and *g*_*random*_ (*r*) is a reference pair distribution function, computed from randomly drawn particles within the convex hull of the nucleosome positions.

### Linker prediction

We have developed a physics-based algorithm to infer linker connections between nucleosomes in tomograms. The algorithm utilizes the positions and orientations of nucleosomes obtained after M-refinement to compute the energy required for linker formation, describing the DNA with the wormlike chain (WLC) model. Based on these energy calculations, probabilities are determined for all possible nucleosome pairings and their respective linker configurations. Linkers are then assigned iteratively, beginning with the configurations of the highest probability. The subsequent sections describe the theory, implementation details, and algorithm validation.

### Theory

Nucleosomes are connected by double-stranded (ds)DNA linkers. Single-molecule elasticity measurements have shown that the entropic elasticity of dsDNA in aqueous solutions is accurately described by the WLC model with a fixed contour length ^56-58^. Deviations of the chain from its straight configuration result in a bending energy *U*_*bend*_^59^. To second order, we approximate the DNA bending energy as ^60^.

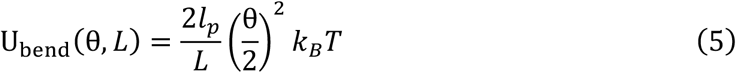

where *l*_*p*_ ≈ 50 nm is the persistence length of DNA ^61^ at physiological conditions, *L* its length, *θ* its bending angle, *k*_*B*_ the Boltzmann constant, and *T* the temperature. We use equation (5) to estimate the energy required to bend a segment of DNA into a linker between the selected arms of a pair of nucleosomes.

### Implementation

Starting from the nucleosomes’ coordinates and orientations, we want to determine the most likely trajectory of the DNA that connects two nucleosomes. From the map obtained from STA (Fig 1e-g), the approximate direction of the two linker arms in the nucleosome is determined (See Supplementary Fig 5a). We denote the linkers of the nucleosome *i* as: Linker1_*i*_, and Linker2_*i*_. There are 4 possible ways Γ to connect a pair of nucleosomes (*i, j*), that is: Γ ∈ [(Linker1_*i*_: Linker), (Linker1_*i*_: Linker2_*j*_), (Linker2_*i*_: Linker1_*j*_), (Linker2_*i*_: Linker2_*j*_)]. For each pair of nucleosomes *i*,*j*, and each combination of k ∈ Γ, we determine the probability of this connection as:

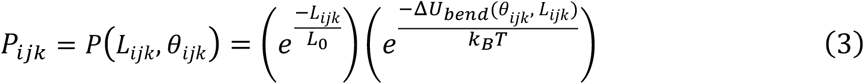

Note that the length *L*_*ijk*_ and the angle *θ*_*ijk*_, depend on the case Γ (see Supplementary Movie 1). In Equation (3), we penalize long linker lengths as our uncertainty increases. We set *L*_0_ = 15 nm as an average value of the expected linker length since DNA linkers range from ∼20-90 bp and vary among different species, tissues, and even fluctuate within a single cellular genome _62_.

To determine *P*_*ijk*_, we computed *L*_*ijk*_ and *θ*_*ijk*_ for a given connection k∈ Γ from the tangent vectors ***T***_*i*_, ***T***_*j*_ and the connecting vector ***D***_*ij*_ (See Supplementary Fig 5a).

### Calculation of the probabilities *P*_*ijk*_

To identify the most likely nucleosome connections, we computed the probabilities *P*_*ijk*_, for all pairwise combinations and cases. To reduce the computational cost, the particle list was pre-processed using cryoCAT ^31^. Using the previously described chain tracing algorithm ^63^, nucleosomes that are spatially close to each other were grouped into chains. For the chain tracing, nucleosome coordinates were shifted to the position of bp 1 and bp 147 (points a_i_ and a_j_ in Supplementary Fig 5a), generating two coordinate sets per nucleosome. Tracing decisions were based solely on the distance between a_i_ and a_j_, with a maximum distance of 20 nm. For each traced cluster *M*, pairwise probability “matrix” *P*_*M*_ ∈ R^N×N×4^ was calculated (See Fig 3b and Supplementary Fig 5c), where |M|=N is the number of particles in the traced cluster, and the third dimension of the matrix corresponds to the four possible configurations, |Γ|=4.

### Assignment of linker connections

From the matrix *P*, we iteratively established the connectivity between particles. First, we defined:

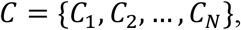

where *C*_i_ represents the set of nucleosomes that are connected to nucleosome *i* and the configuration Γ of the connection. Each particle can have at most one connection at linker 1 and one at linker 2.

For the iterative assignment, we identified the position (*i*,*j*,*k*) with the highest probability *P*_*ijk*_ = *max*(*P*) and checked if the connection (i↔j) was possible based on the topological constraints mentioned above. If the connection is valid, we added (i↔j) to *C*_*i*_, and (j↔i) to *C*_*i*_.

Next, we set *P*_*ij*0_ = *P*_*ij*1_ = *P*_*ij*2_ = *P*_*ij*3_ = 0 and *P*_*ji*0_ = *P*_*ji*1_ = *P*_*ji*2_ = *P*_*jj*3_ = 0, thereby removing the pair of nucleosomes from future iterations. This process continued until no more connections were possible, imposing a probability threshold, *P*_*ijk*_ > 0.1. At each iteration, it was also checked that there are no closed loops.

The connectivity structure *C* can be represented as a graph *G*(*V. E*) where, V are the nucleosomes (nodes) and E are the established connections (edges). See Fig 3c and Supplementary Fig 5c.

### Validation of the predicted linkers

Using the final connectivity graph *G*, wea estimated the coordinates and orientation of the linker centers. A particle was placed at the geometric center between the predicted connected arms, and the orientation of the linkers was assigned to align with the vector connecting the corresponding linker arms (See Supplementary Fig 5a).

### Subtomogram averaging of the predicted linkers

We performed STA in Relion3.1 ^50^ on the 7163 predicted linkers using the positions and orientations derived from the algorithm (see above) with subtomograms being extracted with Warp ^52^, as before no interpolation was used for the STA. The average of the extracted subtomograms was used as the initial reference to avoid biasing the refinement. As a further test, we grouped the linkers based on their predicted length (Fig 4c) and performed STA on each group (see Supplementary Fig 6), resulting in cylindrical shapes with lengths consistent with the predictions.

### *P*_*max*_/*P*_*second*_ of the predicted linkers

To evaluate the certainty of linker assignments in the connectivity analysis, we recorded the highest probability *P*_*max*_ and the second highest probability *P*_*second*_, at each iteration of the algorithm where a linker was assigned. Specifically, 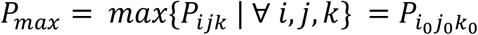 represents the highest probability for the assigned connection between a pair of particles *i* and *j*, reflecting the most confident linkage based on the computed probabilities. On the other hand, 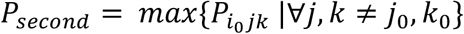 is the second-highest probability value from the same row in the probability matrix, serving as a measure of competing linkage possibilities. A large ratio *P*_*max*_**/***P*_*second*_ indicates a more confident linkage assignment (see Supplementary Fig 7).

### Analysis of the connected nucleosomes

To investigate the structural organization and spatial relationships of connected nucleosomes, we analyzed the spatial distances, and angular relationships between particles in 3D. First, we calculated the **first-neighbor distance (r**_**i**,**i+1**_): as the Euclidean distance between directly connected particles. We then measured the **second-neighbor distance (r**_**i**,**i+2**_) between particles separated by one intermediate nucleosome. **r**_**i**,**i+2**_ is the distance between the center of particles *i* and *i+2* within sorted connected components. Finally, we evaluated the angle (*θ*_i_) between connected triplets of nucleosomes by computing the angle between particles *i, i+1*, and *i+2* in sorted connected components. The angle was determined using the dot product of vectors connecting the particles *i, i+1 and i+1, i+2*.

### All atom model of the connected nucleosome structures

Starting from the STA positions of the nucleosomes, and the predicted linkers, we created all atom models of the nucleosomes, linkers and intrinsically disordered regions (IDRs) of the histone tails. To model the predicted linkers, we adapted the previously established hierarchical chain growth (HCG) method ^64^, which has been used to efficiently sample the conformational space of disordered proteins and single-stranded RNA ^65^. In the present implementation, we generated a library of input fragments by taking the central seven base pairs of 10,000 regularly spaced frames from a previously published molecular dynamics simulation of a 33 base pair B-DNA helix ^66^. At each stitching move, fragments are aligned along backbone heavy atoms of one of two overlapping base pairs. A move is accepted if the alignment RMSD is below a 0.6 Å cut-off. With this, we generated a library of ensembles of 10,000 structures each at a range of possible linker lengths.

For each pair of connected nucleosomes, we then screened the library by aligning three base pair backbones on either end with the respective entry/exit points of the nucleosomal DNA. We considered structures that yield an alignment RMSD below a 2.5 Å cut-off to be suitable candidate structures. For assembly of the full system, we then drew randomly from candidate sub-ensembles at the shortest length that had at least 20 candidate structures until a clash-free combination was found.

In a final step, we added histone tails to the structure by randomly drawing from HCG ensembles and checking for clashes with the rest of the system. We used human histone-tail sequences to match the folded portions included in the template nucleosome. We created the HCG ensembles using a publicly available implementation (https://bio-phys.pages.mpcdf.de/hcg-from-library/)^67^ that draws from a pre-sampled library of dipeptide fragments covering the full sequence space.

### All atom molecular dynamics simulations of the chromatin segment

We performed molecular dynamics simulations of a system of 13 nucleosomes with predicted linkers modelled explicitly using the Gromacs 2024.2 engine ^68^ with the Amber14SB force-field ^69^ augmented with Parmbsc1 parameters ^70^. Systems were solvated with TIP3P water ^71^, 150 mM KCl and 5 mM MgCl_2_ ensuring overall neutrality. We used Mamatkulov-Schwierz parameters for potassium and chloride ions ^72^ and Grotz et al., parameters ^73^ for magnesium ions. After energy minimization, we equilibrated the system in the NVT ensemble at 310 K for 5 ns at a 1fs timestep using Berendsen temperature coupling ^74^. We performed further equilibration in NPT at 1 bar for 20 ns at a 2 fs timestep using C-rescale pressure coupling. We ran the production simulation for 200 ns using velocity-rescale temperature coupling ^75^ and Parrinello-Rahman pressure coupling ^76^. We used frames every 100 ps for analysis.

To illustrate the dynamics of the system, we drew DNA base pair traces every 5 ns starting at 10 ns by aligning the central nucleosome of a trimer with the xy-plane and calculating the center of mass of complementary base pairs across the trajectory. For visualization we rendered the trimer at 70 ns (Fig 5b). Distances r_i,i+2_ and angles θ_I_ were calculated between centers of mass of the 147 base pairs of each nucleosome.

## Acknowledgments

We thank Jenny Sachweh, Desislava Glushkova, Karen Palacio-Rodriguez and Stefan Schaefer for helpful discussions and Stefanie Böhm for critical reading of the manuscript. The Max Planck Computing and Data Facility is acknowledged for computational resources. We thank Sonja Welsch, Mark Linder, Simone Prinz and Susann Kaltwasser (Central Electron Microscopy Facility, MPI of Biophysics) for assistance with cryo-EM sample preparation and data acquisition. We thank Oezkan Yildiz (Structural Biology Unit facility, MPI of Biophysics) and Thomas Hoffman from EMBL for support in high performance computing.

## Authors contributions

Conceptualization: JPK, SCL, JB, MS, BT, ML, GH, MB

Formal analysis: JPK, SCL, JB, TM, MS, BT

Methodology: JPK, SCL, JB, CP, TM, BT

Investigation: JPK, CP

Visualization: JPK, SCL, JB

Funding acquisition: ML, GH, MB

Resources: ML, GH, MB

Software: SCL, TM, BT

Project administration: ML, GH, MB

Supervision: BT, ML, GH, MB

Writing – original draft: JPK, SCL, GH, MB

Writing – review & editing: JPK, SCL, JB, CP, TM, MS, BT, ML, GH, MB

## Data and materials availability

Chromatosome STA maps reported in this paper will be deposited in the EMDB with accession codes XXX and released upon publication.

The raw tilt series and alignment files for the T cell cryo-ET dataset will be deposited on EMPIAR with accession codes XXX and available upon publication.

## References

1. Misteli, T. The Self-Organizing Genome: Principles of Genome Architecture and Function. Cell 183, 28–45 (2020).

2. Luger, K., Mäder, A. W., Richmond, R. K., Sargent, D. F. & Richmond, T. J. Crystal structure of the nucleosome core particle at 2.8 A resolution. Nature 389, 251–260 (1997).

3. Bednar, J. et al. Structure and Dynamics of a 197 bp Nucleosome in Complex with Linker Histone H1. Mol Cell 66, 384-397.e8 (2017).

4. Fyodorov, D. V., Zhou, B. R., Skoultchi, A. I. & Bai, Y. Emerging roles of linker histones in regulating chromatin structure and function. Nat Rev Mol Cell Biol 19, 192–206 (2018).

5. Chen, P., Li, W. & Li, G. Structures and Functions of Chromatin Fibers. Annu Rev Biophys 50, 95–116 (2021).

6. Dombrowski, M., Engeholm, M., Dienemann, C., Dodonova, S. & Cramer, P. Histone H1 binding to nucleosome arrays depends on linker DNA length and trajectory. Nat Struct Mol Biol 29, 493–501 (2022).

7. Song, F. et al. Cryo-EM study of the chromatin fiber reveals a double helix twisted by tetranucleosomal units. Science 344, 376–380 (2014).

8. Schalch, T., Duda, S., Sargent, D. F. & Richmond, T. J. X-ray structure of a tetranucleosome and its implications for the chromatin fibre. Nature 436, 138–141 (2005).

9. Jentink, N., Purnell, C., Kable, B., Swulius, M. T. & Grigoryev, S. A. Cryoelectron tomography reveals the multiplex anatomy of condensed native chromatin and its unfolding by histone citrullination. Mol Cell 83, 3236-3252.e7 (2023).

10. Zhang, M. et al. Angle between DNA linker and nucleosome core particle regulates array compaction revealed by individual-particle cryo-electron tomography. Nat Commun 15, 4395 (2024).

11. Routh, A., Sandin, S. & Rhodes, D. Nucleosome repeat length and linker histone stoichiometry determine chromatin fiber structure. Proc Natl Acad Sci U S A 105, 8872–8877 (2008).

12. Valouev, A. et al. Determinants of nucleosome organization in primary human cells. Nature 474, 516–520 (2011).

13. Lowary, P. T. & Widom, J. New DNA sequence rules for high affinity binding to histone octamer and sequence-directed nucleosome positioning. J Mol Biol 276, 19–42 (1998).

14. Gibson, B. A. et al. Organization of Chromatin by Intrinsic and Regulated Phase Separation. Cell 179, 470-484.e21 (2019).

15. Zhang, M. et al. Molecular organization of the early stages of nucleosome phase separation visualized by cryo-electron tomography. Mol Cell 82, 3000-3014.e9 (2022).

16. Chen, J. K. et al. Nanoscale analysis of human G1 and metaphase chromatin in situ. bioRxiv (2023).

17. Cai, S., Böck, D., Pilhofer, M. & Gan, L. The in situ structures of mono-, di-, and trinucleosomes in human heterochromatin. Mol Biol Cell 29, 2450–2457 (2018).

18. Wang, B. et al. The molecular basis of lamin-specific chromatin interactions. bioRxiv 0 (2024).

19. Hou, Z., Nightingale, F., Zhu, Y., MacGregor-Chatwin, C. & Zhang, P. Structure of native chromatin fibres revealed by Cryo-ET in situ. Nat Commun 14, 6324 (2023).

20. Li, Y., Zhang, H., Li, X., Wu, W. & Zhu, P. Cryo-ET study from in vitro to in vivo revealed a general folding mode of chromatin with two-start helical architecture. Cell Rep 42, 113134 (2023).

21. Zhou, H. et al. Quantitative Spatial Analysis of Chromatin Biomolecular Condensates using Cryo-Electron Tomography. bioRxiv 0 (2024).

22. Kelley, R. et al. Towards community-driven visual proteomics with large-scale cryo-electron tomography ofChlamydomonas reinhardtii. bioRxiv 0 (2024).

23. Majtner, T. et al. cryoTIGER: Deep-Learning Based Tilt Interpolation Generator for Enhanced Reconstruction in Cryo Electron Tomography. bioRxiv 0 (2024).

24. Turoňová, B., Schur, F. K. M., Wan, W. & Briggs, J. A. G. Efficient 3D-CTF correction for cryo-electron tomography using NovaCTF improves subtomogram averaging resolution to 3.4Å. J Struct Biol 199, 187–195 (2017).

25. Cruz-León, S. et al. High-confidence 3D template matching for cryo-electron tomography. Nature Communications 15, 3 (2024).

26. Turoňová, B. GAPStop(TM) - GPU Accelerated Python-base Stopgap for Template Matching [software]. (2024).

27. Wang, S. et al. Linker histone defines structure and self-association behaviour of the 177 bp human chromatosome. Sci Rep 11, 380 (2021).

28. Buchholz, T.-O., Jordan, M., Pigino, G. & Jug, F. Cryo-CARE: Content-Aware Image Restoration for Cryo-Transmission Electron Microscopy Data. 2019 IEEE 16th International Symposium on Biomedical Imaging (ISBI 2019) 100 (2019).

29. Imai, R. et al. Density imaging of heterochromatin in live cells using orientation-independent-DIC microscopy. Mol Biol Cell 28, 3349–3359 (2017).

30. Percus, J. K. & Yevick, G. J. Analysis of Classical Statistical Mechanics by Means of Collective Coordinates. Physical Review 110, 1–13 (1958).

31. Turoňová, B. cryoCAT [software]. (2024).

32. Lamm, L. et al. MemBrain v2: an end-to-end tool for the analysis of membranes in cryo-electron tomography. bioRxiv 16 (2024).

33. Fedulova, A. S. et al. Molecular dynamics simulations of nucleosomes are coming of age. WIREs Computational Molecular Science 14, 0 (2024).

34. Widom, J. Physicochemical studies of the folding of the 100 A nucleosome filament into the 300 A filament. Cation dependence. J Mol Biol 190, 411–424 (1986).

35. Scheffer, M. P., Eltsov, M. & Frangakis, A. S. Evidence for short-range helical order in the 30-nm chromatin fibers of erythrocyte nuclei. Proc Natl Acad Sci U S A 108, 16992–16997 (2011).

36. Bizhanova, A. & Kaufman, P. D. Close to the edge: Heterochromatin at the nucleolar and nuclear peripheries. Biochim Biophys Acta Gene Regul Mech 1864, 194666 (2021).

37. van Steensel, B. & Belmont, A. S. Lamina-Associated Domains: Links with Chromosome Architecture, Heterochromatin, and Gene Repression. Cell 169, 780–791 (2017).

38. Alagna, N. S., Thomas, T. I., Wilson, K. L. & Reddy, K. L. Choreography of lamina-associated domains: structure meets dynamics. FEBS Lett 597, 2806–2822 (2023).

39. Maruyama, Y. & Nishiyama, A. [Mechanism of secretion of ion fluid in exocrine gland cells]. Nihon Rinsho 44, 1500–1503 (1986).

40. Beck, D. B., Oda, H., Shen, S. S. & Reinberg, D. PR-Set7 and H4K20me1: at the crossroads of genome integrity, cell cycle, chromosome condensation, and transcription. Genes Dev 26, 325–337 (2012).

41. Creyghton, M. P. et al. Histone H3K27ac separates active from poised enhancers and predicts developmental state. Proc Natl Acad Sci U S A 107, 21931–21936 (2010).

42. Schaffer, M. et al. Cryo-focused Ion Beam Sample Preparation for Imaging Vitreous Cells by Cryo-electron Tomography. Bio Protoc 5, e1575 (2015).

43. Mastronarde, D.N. Automated electron microscope tomography using robust prediction of specimen movements. Journal of Structural Biology 152, 36–51 (2005).

44. Hagen, W. J. H., Wan, W. & Briggs, J. A. G. Implementation of a cryo-electron tomography tilt-scheme optimized for high resolution subtomogram averaging. J Struct Biol 197, 191–198 (2017).

45. Grant, T. & Grigorieff, N. Measuring the optimal exposure for single particle cryo-EM using a 2.6 Å reconstruction of rotavirus VP6. eLife 4, e06980 (2015).

46. Wan, W. et al. Structure and assembly of the Ebola virus nucleocapsid. Nature 551, 394–397 (2017).

47. Zheng, S. et al. AreTomo: An integrated software package for automated marker-free, motion-corrected cryo-electron tomographic alignment and reconstruction. J Struct Biol X 6, 100068 (2022).

48. Kremer, J. R., Mastronarde, D. N. & McIntosh, J. R. Computer visualization of three-dimensional image data using IMOD. J Struct Biol 116, 71–76 (1996).

49. Turoňová, B. Fourier3D [software]. (2020).

50. Zivanov, J. et al. New tools for automated high-resolution cryo-EM structure determination in RELION-3. Elife 7, (2018).

51. Tegunov, D., Xue, L., Dienemann, C., Cramer, P. & Mahamid, J. Multi-particle cryo-EM refinement with M visualizes ribosome-antibiotic complex at 3.7 Å inside cells. Nature Methods (2020).

52. Tegunov, D. & Cramer, P. Real-time cryo-electron microscopy data preprocessing with Warp. Nat Methods 16, 1146–1152 (2019).

53. Reda, F. et al. in Lecture Notes in Computer Science: Computer Vision – ECCV 2022 250-266 (Springer Nature Switzerland, Cham, 2022).

54. Campello, R. J. G. B., Moulavi, D. & Sander, J. in Lecture Notes in Computer Science: Advances in Knowledge Discovery and Data Mining 160–172 (Springer Berlin Heidelberg, Berlin, Heidelberg, 2013).

55. Barber, C. B., Dobkin, D. P. & Huhdanpaa, H. The quickhull algorithm for convex hulls. ACM Transactions on Mathematical Software 22, 469–483 (1996).

56. Smith, S. B., Finzi, L. & Bustamante, C. Direct mechanical measurements of the elasticity of single DNA molecules by using magnetic beads. Science 258, 1122–1126 (1992).

57. Perkins, T. T., Smith, D. E., Larson, R. G. & Chu, S. Stretching of a single tethered polymer in a uniform flow. Science 268, 83–87 (1995).

58. Bustamante, C., Marko, J. F., Siggia, E. D. & Smith, S. Entropic elasticity of lambda-phage DNA. Science 265, 1599–1600 (1994).

59. Marko, J. F. Stretching must twist DNA. Europhysics Letters (EPL) 38, 183–188 (1997).

60. Marin-Gonzalez, A., Vilhena, J. G., Perez, R. & Moreno-Herrero, F. Understanding the mechanical response of double-stranded DNA and RNA under constant stretching forces using all-atom molecular dynamics. Proc Natl Acad Sci U S A 114, 7049–7054 (2017).

61. Marko, J. F. & Siggia, E. D. Stretching DNA. Macromolecules 28, 8759–8770 (1995).

62. Szerlong, H. J. & Hansen, J. C. Nucleosome distribution and linker DNA: connecting nuclear function to dynamic chromatin structure. Biochem Cell Biol 89, 24–34 (2011).

63. Xing, H. et al. Translation dynamics in human cells visualized at high resolution reveal cancer drug action. Science 381, 70–75 (2023).

64. Pietrek, L. M., Stelzl, L. S. & Hummer, G. Hierarchical Ensembles of Intrinsically Disordered Proteins at Atomic Resolution in Molecular Dynamics Simulations. J Chem Theory Comput 16, 725–737 (2020).

65. Pietrek, L. M., Stelzl, L. S. & Hummer, G. Hierarchical Assembly of Single-Stranded RNA. J Chem Theory Comput 20, 2246–2260 (2024).

66. Cruz-León, S. et al. Twisting DNA by salt. Nucleic Acids Res 50, 5726–5738 (2022).

67. Pietrek, L. M., Stelzl, L. S. & Hummer, G. Structural ensembles of disordered proteins from hierarchical chain growth and simulation. Curr Opin Struct Biol 78, 102501 (2023).

68. Abraham, M. J. et al. GROMACS: High performance molecular simulations through multi-level parallelism from laptops to supercomputers. SoftwareX 1-2, 19–25 (2015).

69. Maier, J. A. et al. ff14SB: Improving the Accuracy of Protein Side Chain and Backbone Parameters from ff99SB. J Chem Theory Comput 11, 3696–3713 (2015).

70. Ivani, I. et al. Parmbsc1: a refined force field for DNA simulations. Nat Methods 13, 55–58 (2016).

71. Jorgensen, W. L., Chandrasekhar, J., Madura, J. D., Impey, R. W. & Klein, M. L. Comparison of simple potential functions for simulating liquid water. The Journal of Chemical Physics 79, 926–935 (1983).

72. Mamatkulov, S. & Schwierz, N. Force fields for monovalent and divalent metal cations in TIP3P water based on thermodynamic and kinetic properties. J Chem Phys 148, 074504 (2018).

73. Grotz, K. K., Cruz-León, S. & Schwierz, N. Optimized Magnesium Force Field Parameters for Biomolecular Simulations with Accurate Solvation, Ion-Binding, and Water-Exchange Properties. J Chem Theory Comput 17, 2530–2540 (2021).

74. Berendsen, H. J. C., Postma, J. P. M., van Gunsteren, W. F., DiNola, A. & Haak, J. R. Molecular dynamics with coupling to an external bath. The Journal of Chemical Physics 81, 3684–3690 (1984).

75. Bussi, G., Donadio, D. & Parrinello, M. Canonical sampling through velocity rescaling. J Chem Phys 126, 014101 (2007).

76. Parrinello, M. & Rahman, A. Polymorphic transitions in single crystals: A new molecular dynamics method. Journal of Applied Physics 52, 7182–7190 (1981).

