## Supplementary Information for "Molecular architecture of heterochromatin at the nuclear periphery of primary human cells"

**The PDF file includes:**

Supplementary Figures 1 to 8

Supplementary Table 1

**Other Supplementary Information for this manuscript include the following:**

Supplementary Movies 1 to 3

**Supplementary Figure 1.**

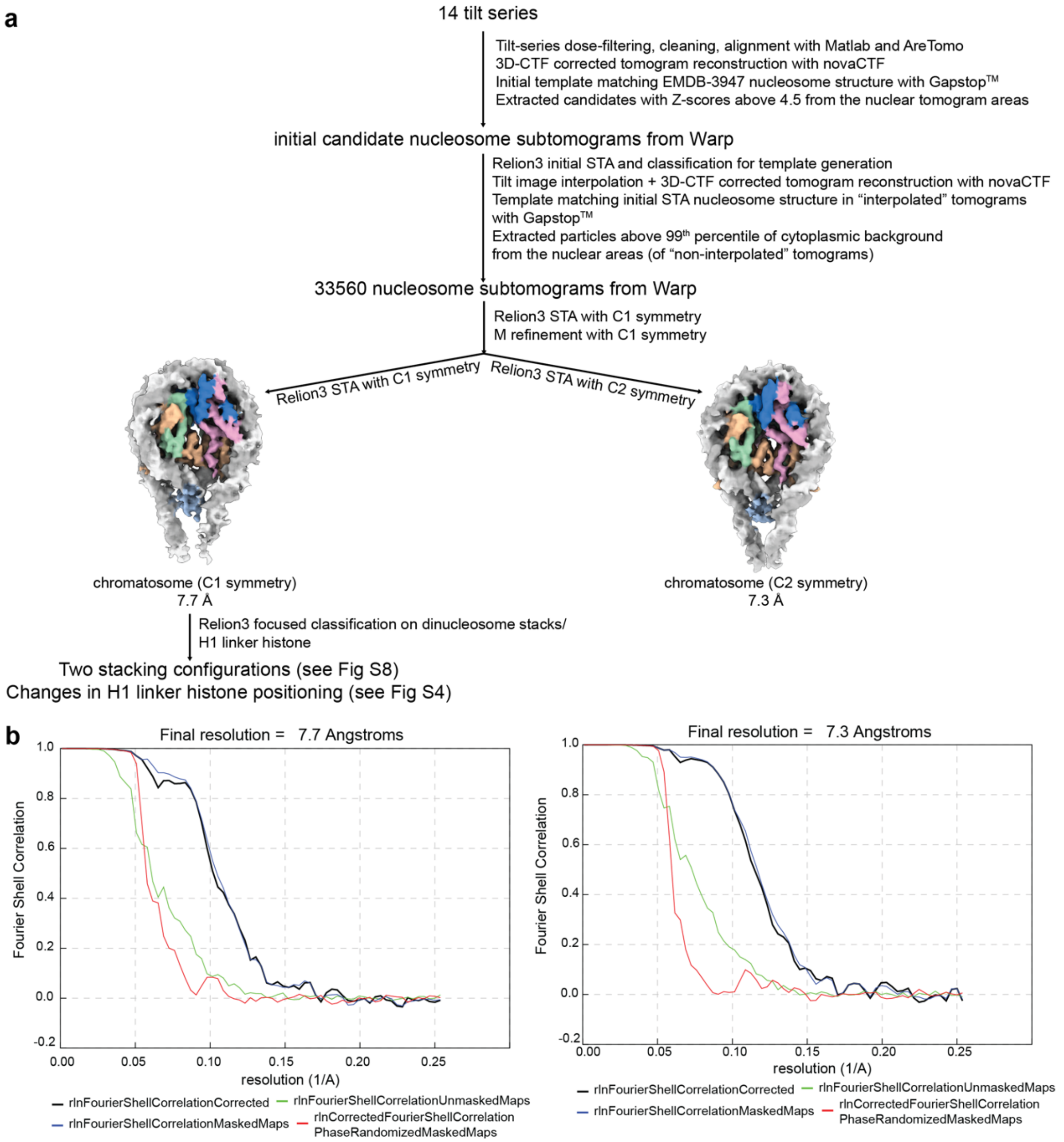


**Image processing workflow for the chromatosomes. a** Workflow describing the steps taken from raw tilt series to final chromatosome STA structures. **b** Fourier shell correlation curves from Relion 3.1 [50] after M refinement [51] for structures with C1 or C2 symmetry imposed.

**Supplementary Figure 2.**

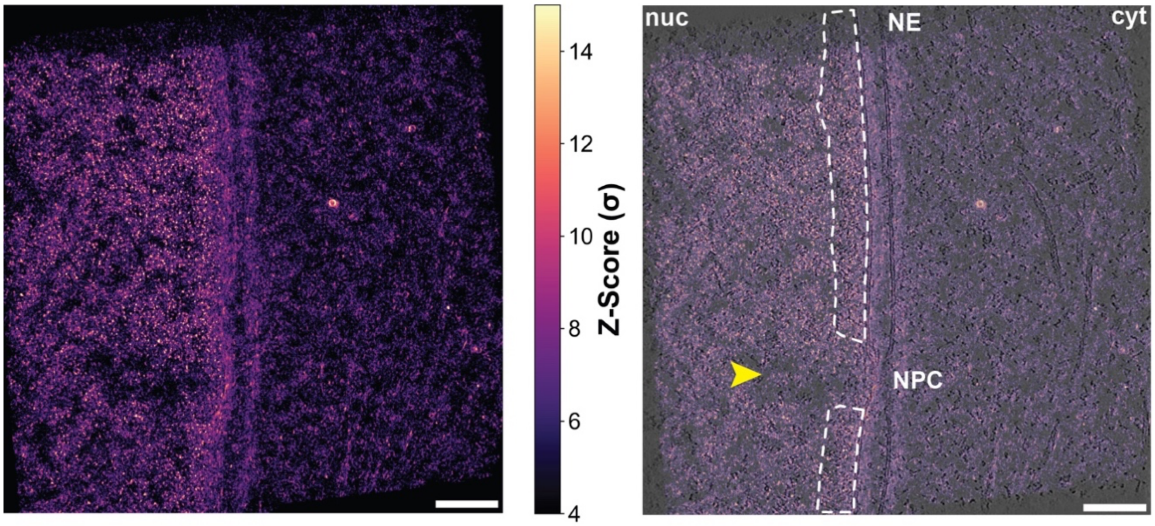


**Template matching nucleosomes with GAPSTOP$^{TM}$ [26].** Constrained cross-correlation (CCC) volume (transformed to Z-scores) from template matching nucleosome structure on an exemplary tomogram, shown as a maximum intensity projection and color-coded by Z-score. The number of and intensity of the peaks in the nuclear region of the tomogram is much higher as can be seen in the overlay of the CCC volume with a tomogram slice. Chromatin-free space is indicated with a yellow arrow, a dense layer of chromatin underlying the nuclear envelope is framed white.

**Supplementary Figure 3.**

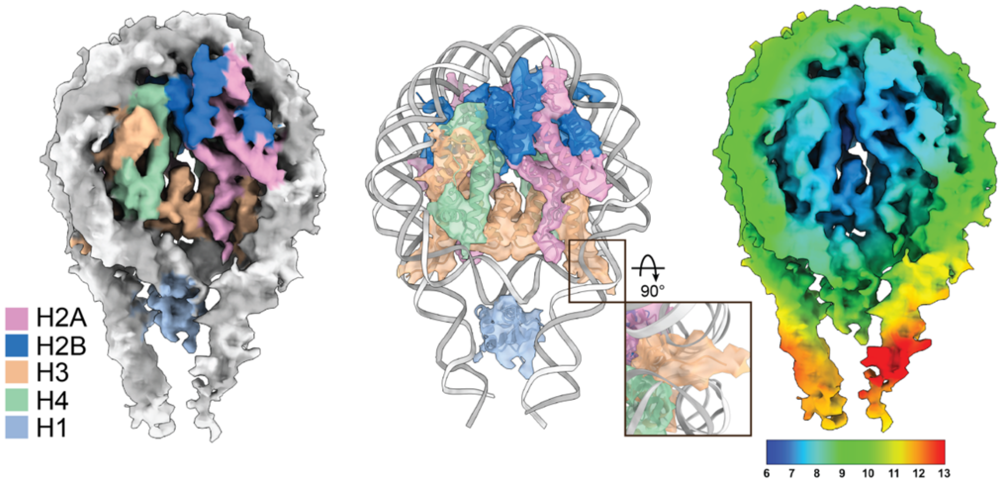


**STA of the chromatosome with C1 symmetry.** Color-coded STA map by DNA strands (white and gray) and histone proteins H2A (pink), H2B (blue), H3 (amber), H4 (green) and H1 (light blue). The atomic model of the canonical chromatosome (PDB: 7dbp [27]) was fitted into our map and is shown in the middle panel. On the right, a color-coded local resolution STA map of the human chromatosome (from high resolution in blue to low resolution in red, in Å).

**Supplementary Figure 4.**

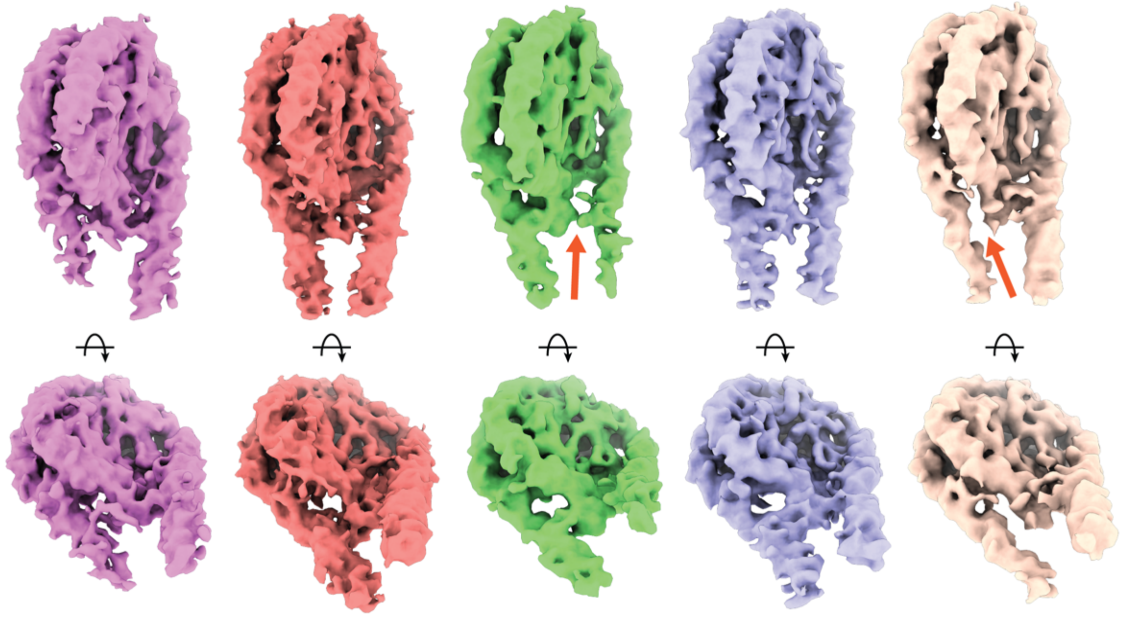

**Focused 3D classification around H1 linker histone area reveals shifting H1 histone.** 3D classification in Relion 3.1 [50] identified nucleosome classes in which H1 was shifted to either DNA linker, characteristic for off dyad binding (highlighted with orange arrows).

**Supplementary Figure 5.**

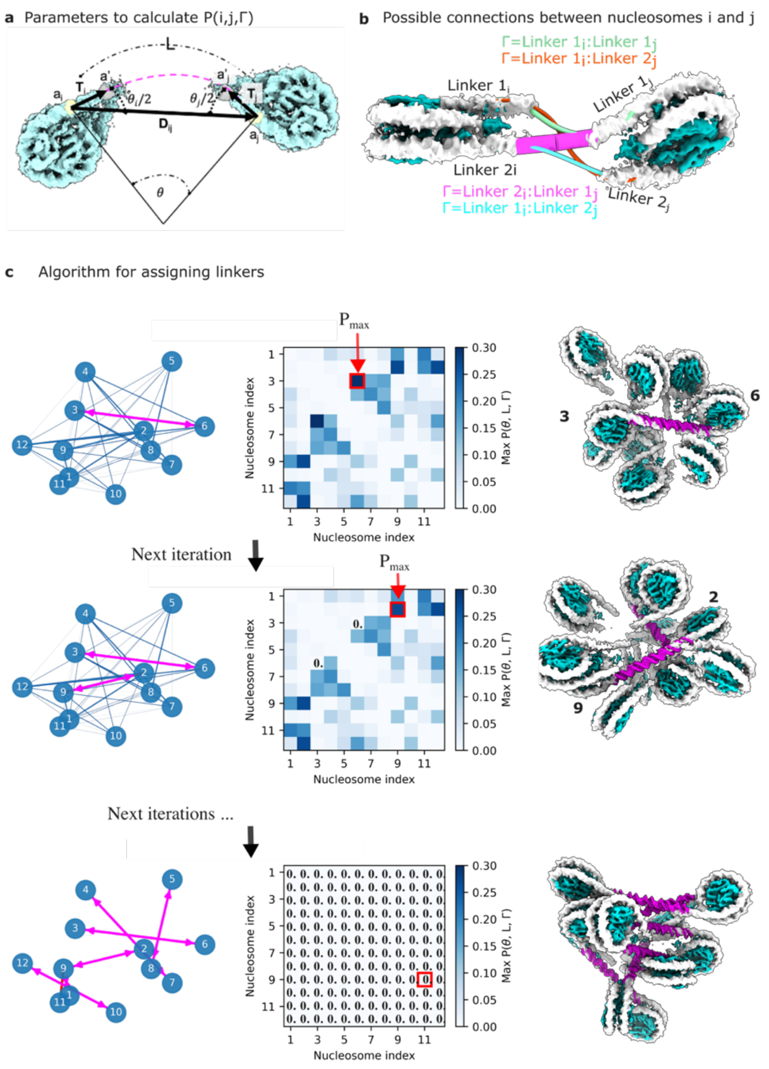


**Visualization of the linker assignment. a** Scheme for determining the parameters to for the probability of linkers. The probability of a potential linker (pink dashed line) to exist is calculated from the bending angle, $\theta$, and the arc length $L$. For each pair of nucleosomes $i,j$, and each combination Γ of connections between the arms, the length L is determined as the arc length measure from the center of bp 1 or bp 147 in the nucleosome (points $a_i$, and $a_j$ marked with yellow spheres). The bending angle is determined as $\theta = \frac{\theta_i+\theta_j}{2}$. **b** Set of possible connections Γ between a pair of nucleosomes (*i, j*). **c** Algorithm for assigning linker connections. The columns show the graph of the nucleosomes (left), the probability matrix P (middle) obtained as described in Methods, and the visualization of the nucleosome position and the predicted linkers (right). The linkers are assigned iteratively, starting with the configurations with the highest probability $P_{max}$. In the example shown the linker (3↔6) is created in the first iteration. For the next iteration, we set $P_{36k} = P_{63k} = 0, \forall\, k \in [0,1,2,3]$, and now $P_{max}$ creates the connection (2↔9). The process continues until no further connections are possible with a threshold $P_{ijk} > 0.1$. Note that the indices of the particles is arbitrary.

**Supplementary Figure 6.**

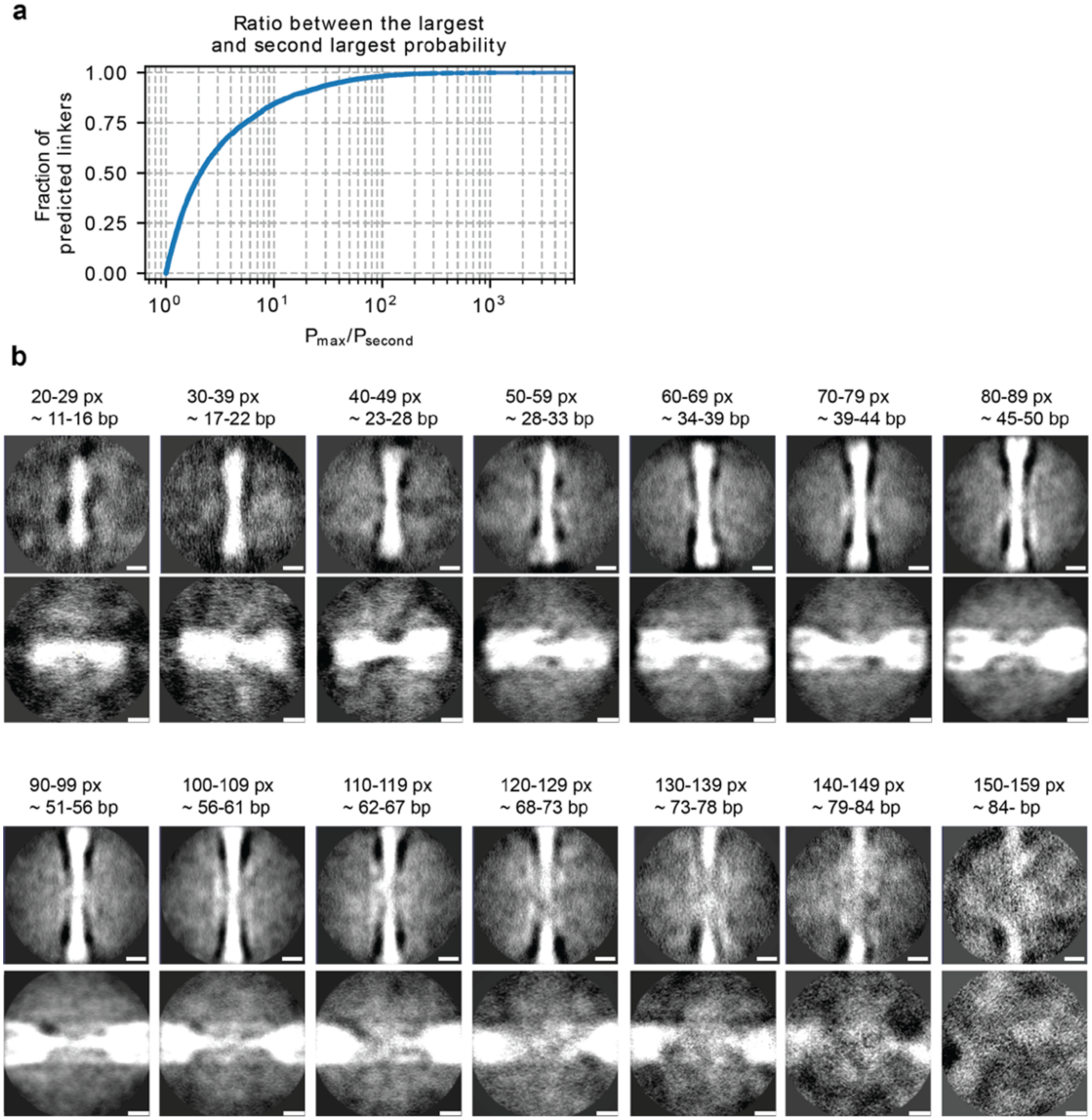


**Predicted DNA linker validation through $P_{max}/P_{second}$ and STA. a** Cumulative distribution of the ratio between the highest probability $P_{max}$ and the second highest probability $P_{second}$, for all the assigned linkers (see Methods). This ratio quantifies competing linkage possibilities, with larger values indicating higher confidence in the linkage assignment. **b** The DNA linkers were grouped by predicted length and subjected to separate STA refinements in Relion 3.1 [50]. YZ and XZ slices through each linker DNA average are shown with the predicted length matching well to increasing length of the densities.

**Supplementary Figure 7.**

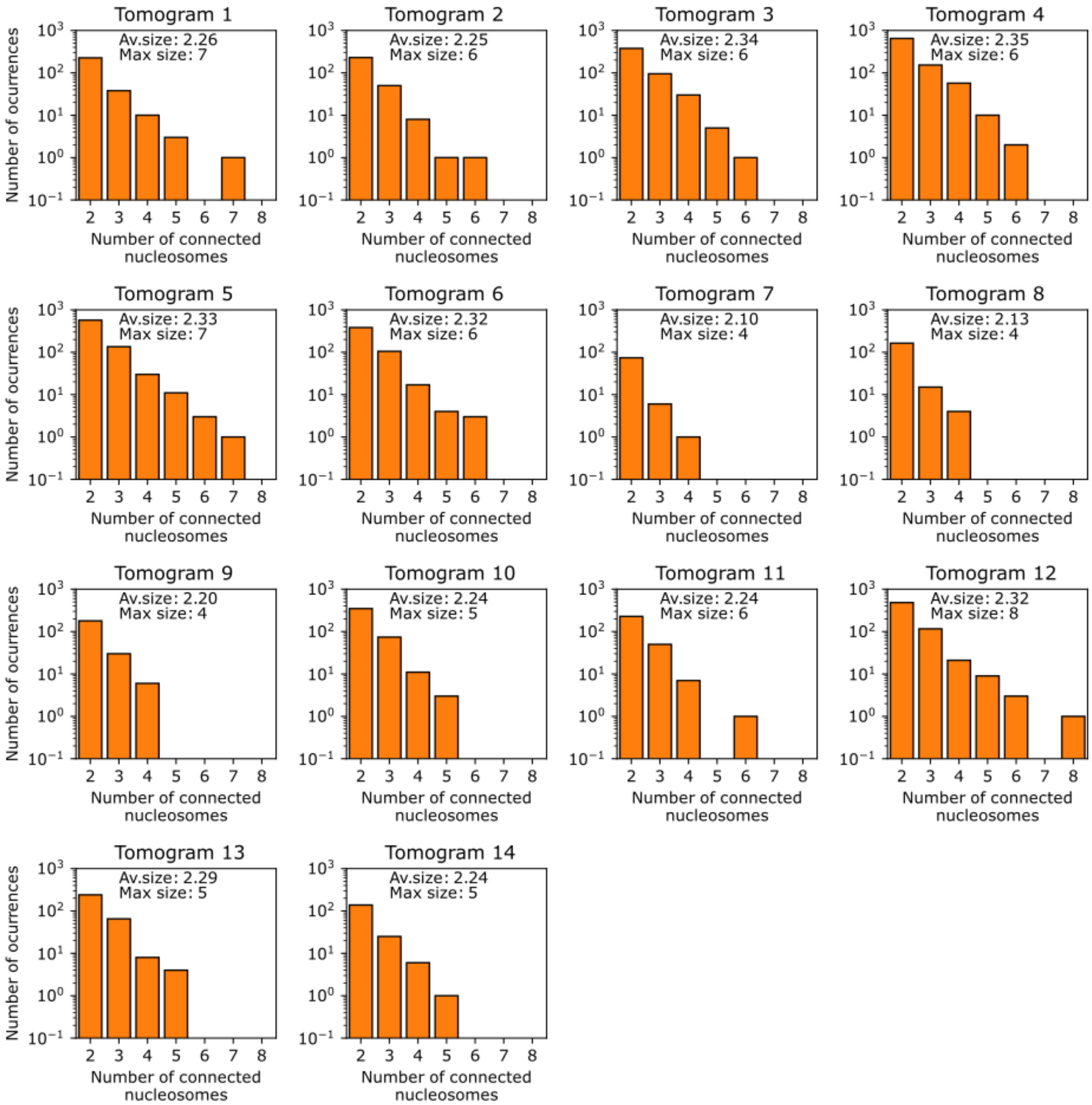


**Histograms of the number of connected nucleosomes per tomogram**. Distribution of the number of connected nucleosomes per tomogram after the assignment of linker connections. The inset text shows the maximum number of connected nucleosomes per tomogram, as well as the average size.

**Supplementary Figure 8.**

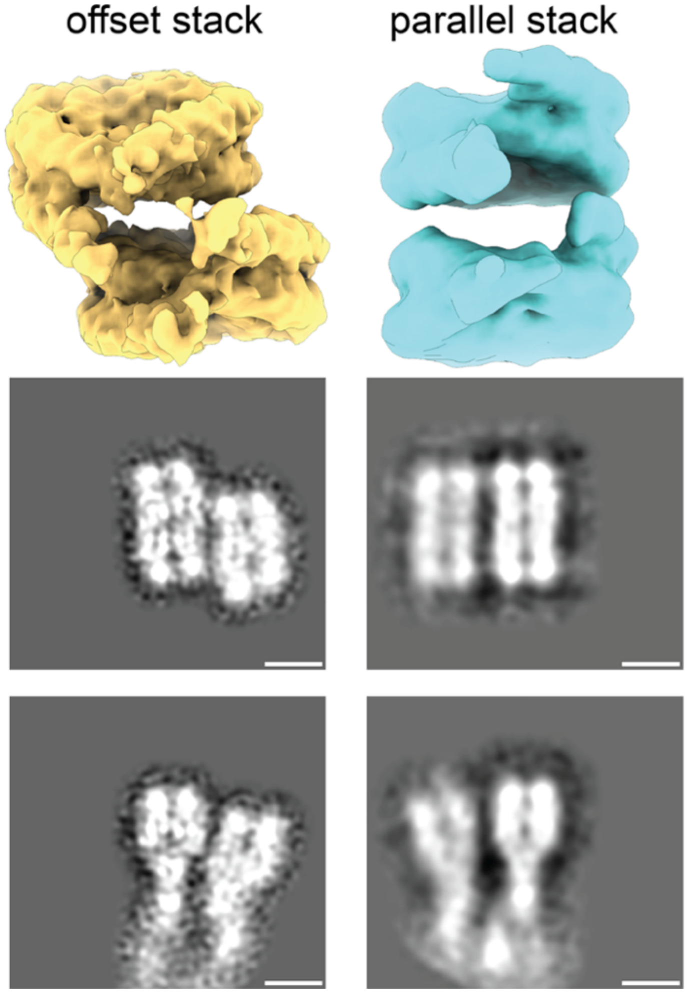


**3D classification of stacked nucleosomes reveals two stacking configurations**. 3D classification in Relion 3.1 [50] found that ~18% of all nucleosomes were in one of two stacking configurations. Stacking configuration 1 shows the particles stacked with an offset whereas stacking configuration 2 shows the particles stacked directly on top of each other. The center-to-center distance between the stacked nucleosomes was ~6.5 nm. Scale bars: 5 nm.

**Supplementary Table 1: cryo-ET data acquisition parameters and STA map information.**

| | | |
|---|---|---|
| **Microscope** | Titan Krios G4 | |
| **Voltage (kV)** | 300 | |
| **Camera** | Falcon 4 | |
| **Magnification** | 64000 | |
| **Pixel size (Å/px)** | 1.971 | |
| **Targeted total electron dose ($e^{-}/Å^{2}$)** | 135 | |
| **Targeted defocus range (µm)** | -1.75 – 4.25 | |
| **Automation software** | SerialEM | |
| **Tomograms used for. STA/TM** | 14 | |
| **Initial # of nucleosomes** | 33560 | |
| **Map type** | Chromatosome with C1 symmetry | Chromatosome with C2 symmetry |
| **Final # of particles** | 33560 | 33560 |
| **Resolution (Å) (FSC 0.143)** | 7.7 | 7.3 |

**Supplementary Movie 1.**

Possible connections between nucleosomes.

**Supplementary Movie 2.**

Assignments of the elements in the connectivity matrix: Example of the connectivity explaining the procedure and showing that STA averages already 'draw' the linkers.

**Supplementary Movie 3.**

MD simulation of the 13-chromatosomes.
